# *Drosophila* PDGF/VEGF signaling from muscles to hepatocyte-like cells protects against obesity

**DOI:** 10.1101/2019.12.23.887059

**Authors:** Arpan C. Ghosh, Sudhir G. Tattikota, Yifang Liu, Aram Comjean, Yanhui Hu, Victor Barrera, Shannan J. Ho Sui, Norbert Perrimon

**Affiliations:** Department of Genetics, Blavatnik Institute, Harvard Medical School, Boston, USA; Howard Hughes Medical Institute, Boston, USA; Harvard T.H. Chan Bioinformatics Core, Boston, USA

**Keywords:** *Drosophila*, VEGF, PDGF, Pvf, PvR, mTOR, obesity, myokine, hepatocyte, liver, oenocytes, adipose tissue, lipid metabolism, snRNA-Seq

## Abstract

PDGF/VEGF ligands regulate a plethora of biological processes in multicellular organisms via autocrine, paracrine and endocrine mechanisms. Here, we investigated organ-specific roles of *Drosophila* PDGF/VEGF-like factors (Pvfs). We combine genetic approaches and single-nuclei sequencing to demonstrate that muscle-derived Pvf1 signals to the *Drosophila* hepatocyte-like cells/oenocytes to suppress lipid synthesis by activating the Pi3K/Akt1/mTOR signaling cascade in the oenocytes. Additionally, we show that this signaling axis regulates the rapid expansion of adipose tissue lipid stores observed in newly eclosed flies. Flies emerge after pupation with limited adipose tissue lipid stores and lipid levels are progressively restored via lipid synthesis. We find that *pvf1* expression in the adult muscle increase rapidly during this stage and that muscle-to-oenocyte Pvf1 signaling inhibits restoration of adipose tissue lipid stores as the process reaches completion. Our findings provide the first evidence in a metazoan of a PDGF/VEGF ligand acting as a myokine that regulates systemic lipid homeostasis by activating mTOR in hepatocyte-like cells.

**Highlights:**

- Muscle specific Pvf1 protects mature adult flies from obesity
- Single-nuclei RNA sequencing reveals that PvR, the receptor for Pvf1, is highly expressed in the *Drosophila* hepatocyte-like cells/oenocytes.
- PvR is required specifically in oenocytes to protect adult flies from obesity
- Muscle-to-oenocyte Pvf1 signaling activates PvR/Pi3K/Akt1/mTOR in the oenocytes to suppress lipid synthesis
- Muscle-derived Pvf1 helps terminate the rapid expansion of adipose tissue lipid stores in newly eclosed flies

## Introduction

Specialized organ systems compartmentalize core metabolic responses such as nutrient uptake, nutrient storage, feeding behavior and locomotion in multicellular organisms. In order to link systemic metabolic status to appropriate physiological responses, information on local metabolic events within each of these organs must be integrated. Inter-organ communication factors play an important role in mediating this process of systemic metabolic integration. For instance, classical metabolic hormones such as insulin and glucagon released from the pancreas, as well as leptin released from the adipose tissue, can act on multiple peripheral and central nervous system targets. While insulin and glucagon define anabolic and catabolic states of an animal, leptin limits food intake in response to adequate energy stores in the adipose tissue (Ahima et al., 1996; Moore and Cooper, 1991; Schade et al., 1979; Tartaglia et al., 1995). In addition to these classic hormones, a number of peptides have been identified that can mediate inter-organ communication axes. For example, adiponectin, adipisin and asprosin are molecules that are released from the adipose tissue that signal to distant tissues such as muscle/liver and pancreas (Lo et al., 2014; Romere et al., 2016; Yamauchi et al., 2014). IGF1, angiotensin and IGFBPs are released from the liver and signal to multiple distant organs including adipose tissue, muscle and kidney (Boucher et al., 2012; Clemmons, 2007; Droujinine and Perrimon, 2016). The role of the skeletal muscle as an endocrine organ has gained special interest lately, primarily due to the beneficial effects of having healthy active muscles towards ameliorating or preventing the pathophysiology of a number of disorders and diseases (Benatti and Pedersen, 2015; Boström et al., 2012; So et al., 2014). A number of skeletal muscle-derived signaling factors (“myokines”), including Irisin, Myostatin, IL6, Myonectin and FGF21, have recently been characterized and they signal both locally, and to distant tissues, to control processes as diverse as muscle growth and browning of fat (Boström et al., 2012; Fisher and Maratos-Flier, 2016; McPherron, 2010; Pedersen and Febbraio, 2012; Seldin et al., 2012). Nevertheless, conservative estimates, based on bioinformatic analysis of the skeletal muscle transcriptome and proteomics studies, indicate that skeletal muscles are capable of secreting more than 200 myokines and the vast majority of these proteins are yet to be characterized (Pedersen and Febbraio, 2012). The vertebrate PDGF/VEGF signaling ligands are also secreted from the skeletal muscles; however, the biological roles of muscle-derived PDGF/VEGF ligands remain unknown (Catoire et al., 2014; Henningsen et al., 2010; Raschke et al., 2013).

The PDGF/VEGF family of signaling ligands have co-evolved with multicellularity and are proposed to signal in the context of organisms with tissue-level organization (Holmes and Zachary, 2005). VEGF family ligands, including VEGF-A and VEGF-B, influence metabolic responses such as adiposity, insulin resistance and browning of fat (Elias et al., 2012; Hagberg et al., 2012; Lu et al., 2012; Robciuc et al., 2016; Sun et al., 2012; Sung et al., 2013; L. E. Wu et al., 2014). However, mechanisms by which these molecules regulate lipid metabolism are complex and multiple context dependent and confounding models of their action have been proposed (Elias et al., 2012; Hagberg et al., 2012; Lu et al., 2012; Robciuc et al., 2016; Sun et al., 2012; Sung et al., 2013; L. E. Wu et al., 2014). Nevertheless, in all of these studies the effect of VEGFs on obesity and insulin resistance have been attributed to their roles in modulating tissue micro-vascularization and endothelial cell biology. Similarly, PDGF signaling ligands PDGF-BB and -CC have also been implicated in adipose tissue expansion, glucose metabolism and thermogenesis in beige fat by influencing remodeling of tissue vascularization (Onogi et al., 2017; Seki et al., 2016). VEGFs and PDGFs are well known regulators of vascularization and endothelial cell biology, and tissue vascularization definitely plays an important role in adipose tissue health, inflammation and ultimately insulin resistance. Nevertheless, signaling to non-endothelial cell types by PDGF/VEGF ligands could play equally important roles in regulating obesity and insulin resistance.

In *Drosophila*, the PDGF/VEGF pathway ligands are encoded by three genes, *pvf1*, *pvf2* and *pvf3* (Cho et al., 2002; Duchek et al., 2001). Similar to vertebrate PDGFs/VEGFs, these molecules signal through a Receptor-Tyrosine Kinase (RTK) encoded by *pvr*. Once bound to the receptor, Pvfs primarily activate the Ras/Raf/ERK intracellular cascade (Duchek et al., 2001; Heino et al., 2001).

Phylogenetic analysis of the Pvfs show that Pvf1 is closely related to both VEGFs and PDGFs and most likely plays the dual role of representing both these divergent signaling pathways in the fly (Holmes and Zachary, 2005). Pvf2 and Pvf3 are more ancestral forms of the protein and are phylogenetically distinct with Pvf2 lacking the conserved cysteine necessary for forming the cysteine knot structure characteristic of this family of ligands (Holmes and Zachary, 2005; Kasap, 2006).

Core metabolic organs and signaling mechanisms that regulate metabolic homeostasis are also highly conserved between *Drosophila* and vertebrates (Droujinine and Perrimon, 2016). Metabolic organ systems that are functionally and structurally analogous to the vertebrate adipose tissue, muscle, intestine, and liver exist in *Drosophila* (Gutierrez et al., 2007; Leopold and Perrimon, 2007). The *Drosophila* adipose tissue (also called the fat body) functions as the primary site for lipid storage. The adipose tissue is also the key site for sensing the nutrient status of the animal and coupling it to systemic growth, metabolism and feeding behavior (Colombani et al., 2003; Grönke et al., 2007). The roles of the liver are shared between the adipose tissue and specialized hepatocyte-like cells called the oenocytes, with the adipose tissue being the primary site of glycogen storage and the oenocytes playing the roles of lipid mobilization and synthesis of specialized lipid molecules (Gutierrez et al., 2007; Makki et al., 2014; Storelli et al., 2019). However, much remains to be learned about the biological roles of the specialized organ systems in *Drosophila* and how they relate to organ systems in vertebrates.

A number of hormonal signals emanating from the *Drosophila* adipose tissue, gut and muscle have been characterized and shown to play roles as diverse as regulation of insulin and glucagon (AKH in *Drosophila*) release, nutrient uptake in the gut and mitochondrial metabolism (Chng et al., 2014; Droujinine and Perrimon, 2016; Ghosh and O’Connor, 2014; Rajan and Perrimon, 2012; Song et al., 2017). Components of core metabolic pathways such as the mTOR signaling pathway are also highly conserved in the fly and much has been learned about the fundamental principles of the regulation of mTOR and its roles in growth and aging from studies in *Drosophila* (Antikainen et al., 2017; Colombani et al., 2003; Kim et al., 2008; Piper and Partridge, 2018). *Drosophila* Pvfs have been primarily studied in the context of their roles in embryonic development, cell motility and specification of immune cells (Duchek et al., 2001; Ishimaru et al., 2004; Rosin et al., 2004). However, the role of these signaling peptides in metabolism remains less explored.

Here, we identify the *Drosophila* PDGF/VEGF ortholog, Pvf1, as a tubular muscle derived signaling factor (myokine) that regulates systemic lipid stores by inhibiting lipid synthesis. Additionally, by subjecting the metabolically active organ systems that reside in *Drosophila* abdomen (muscle, oenocytes and adipose tissue) to single-nuclei RNA sequencing (snRNA-Seq), we identify the adult oenocytes to be one of the primary targets of Pvf signaling. We provide further evidence that muscle-derived Pvf1 signals to the oenocytes where it activates the PvR/Pi3K/Akt1/mTOR signaling pathway that in turn inhibits lipid synthesis in the organism. We also find that muscle-derived *pvf1* plays an important role in regulating the restoration of adipose tissue lipid stores in newly eclosed flies. Flies eclose from their pupal case with limited adipose tissue lipid stores. Post eclosion they enter a developmental stage that is marked by increased lipid synthesis needed for restoration of the adipose tissue lipid stores. *pvf1* expression in the muscle increases rapidly during this stage and premature expression of *pvf1* in the muscle interferes with the process of lipid restoration. Taken together our study indicates that muscle-Pvf1 serves as an inhibitory signal for suppressing de-novo lipid synthesis once the adult adipose tissue has accumulated sufficient lipid stores at the end of the adipose tissue lipid-restoration phase. Our snRNA-seq data of the adult abdominal region that includes adipose tissue, oenocytes, and abdominal muscles will also serve as an invaluable resource to further understand the biology of these organs. To facilitate visualization of this rich resource of gene expression profiles, we have created a searchable webtool where users can mine and explore the data (https://www.flyrnai.org/scRNA/abdomen/).

## Results

### Muscle-specific loss of Pvf1 leads to increased lipid accumulation in the adipose tissue and oenocytes

To investigate tissue specific roles of Pvfs in regulating lipid homeostasis, we knocked down *pvf1*, *pvf2* and *pvf3* in various metabolically active tissues in the adult male fly using temperature inducible drivers and looked for effects on the adiposity of the fly. Strikingly, knockdown of *pvf1* in the adult muscle (*mus^ts^>pvf1-i*), but not in other organs such as the gut, adipose tissue or oenocytes, was associated with a severe obesity phenotype characterized by increased lipid droplet size in the adipose tissue cells (Fig. 1-A,B and Fig. S1-A). Additionally, the experimental animals showed increased accumulation of lipid droplets in the oenocytes which are normally devoid of or have very few lipid droplets (Fig. 1-A,B and Fig. S1-A). *mus^ts^>pvf1-i* flies also showed significantly higher levels of whole animal triacylglycerol (TAG) content than control flies (Fig. 1-C). The increase in total TAG content was observed for two independent RNAi lines against *pvf1* (Fig. 1-C). The increase in TAG content of *mus^ts^>pvf1-i* flies was more pronounced when flies were challenged with a mildly high sugar diet (15% w/v added sugar to our standard food) (Fig. S1-B). Note that we used this food condition for all our experiments unless mentioned otherwise. Knocking down either *pvf2* or *pvf3* in the adult muscle did not affect total TAG content of the animals, further demonstrating that the phenotype is specifically associated with muscle-specific loss of Pvf1 (Fig. 1-D). When fed with ^14^C-U-Glucose for 2 days, *mus^ts^>pvf1-i* and control flies showed comparable incorporation of ^14^C in whole fly homogenates (Fig. S1-C), indicating that *mus^ts^>pvf1-i* flies do not eat more than control animals and that the obesity phenotype observed is most likely caused by metabolic defects.

**Figure 1:**
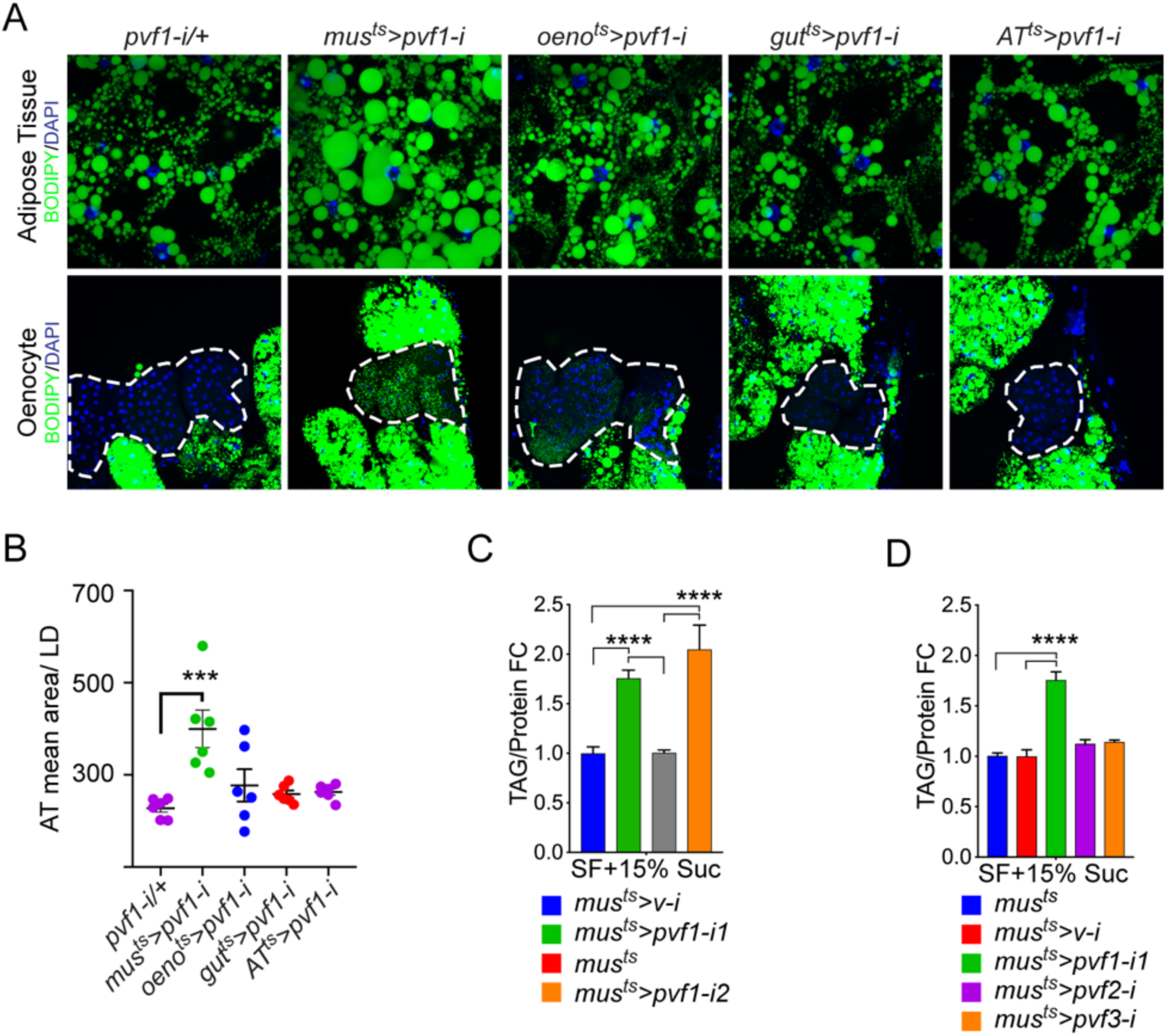
A muscle-to-oenocyte Pvf1 signaling axis protects against obesity (A) BODIPY staining showing neutral lipid accumulation in the adult male adipose tissue (AT) and oenocytes (dorsal abdominal cuticle) of flies in which *pvf1* was knocked down using an RNAi transgene (VDRC: kk102699) specifically in the adult: muscle (*mus^ts^=muscle-Gal4^Gal80ts^*), oenocytes (*oeno^ts^=oenocyte-Gal4^Gal80ts^*), gut (*gut^ts^=gut-Gal4^Gal80ts^*) and adipose tissue (*AT^ts^= AdiposeTissue-Gal4^Gal80ts^*). The RNAi transgene alone is shown as a control. Similar results were observed using a different RNAi line (NIG: 7103R-2, data not shown). (B) Mean lipid droplet size (≥10 microns in diameter) in the adipose tissue of flies shown in Fig. 1A. Only muscle specific loss of *pvf1* show a significant increase (*p<0.001*) compared to the control genotype, One-Way ANOVA followed by Tukey’s HSD test. (C) Triacylglycerol (TAG) assay showing TAG/protein ratio in adult males in which *pvf1* was knocked down specifically in the muscle. Knocking down *pvf1* using two independent RNAi lines (*pvf1-i1*::VDRC kk102699 and *pvf1-i2*::NIG 7103R-2) leads to a significant increase in total TAG content of the flies. (*mus^ts^>v-i=muscle-Gal4^Gal80ts^>vermilion-RNAi*). (SF+15% Suc = standard lab food supplemented with 15% sucrose w/w). n=6, One-Way ANOVA followed by Tukey’s HSD test. (D) Total TAG content of adult males with adult muscle-specific (*mus^ts^*) knockdown of *pvf1*, *pvf2* and *pvf3.* n=6, One-Way ANOVA followed by Tukey’s HSD test.

To verify the presence and distribution of Pvf1 protein in adult muscles we immunostained the indirect flight muscles (IFMs) and leg muscles using an anti-Pvf1 antibody (Duchek et al., 2001; Rosin et al., 2004). Pvf1 is abundantly present in the striated tubular leg muscles (Fig. S1-D) where it is stored between the individual myofibrils (Fig. S1-D1*’*, D3*’*) and is more concentrated at both the M and Z discs (Fig. S1-D3,3*’*,3”). To verify the specificity of the signal we immunostained the leg muscles of *mus^ts^>pvf1-i* flies with anti-Pvf1 antibody and observed a strong reduction in Pvf1 protein level (Fig. S1-D2’,D4,D4’). Interestingly, the IFMs did not show any staining for Pvf1 indicating that the protein is primarily stored in the striated tubular muscles in the fly.

### Single-nuclei RNA-sequencing (snRNA-Seq) identifies PvR RTK signaling pathway enriched in Oenocytes

Single-nuclei sequencing presents unprecedented access to the transcriptomes of cell types residing in complex tissue structures or organs that are difficult to dissect and segregate (Birnbaum, 2018; Chen et al., 2018; Kulkarni et al., 2019; H. Wu et al., 2019). We took advantage of this tool to understand the transcriptomics of tissue types residing in the adult abdominal cuticle that harbors several metabolically active tissues such as the fat body, abdominal-muscles, and oenocytes, which are functionally analogous to adipose tissue, skeletal muscle, and liver, respectively, in higher vertebrates (Droujinine and Perrimon, 2016; Musselman and Kühnlein, 2018). To delineate the patterns of gene expression in each of these tissues, we dissected and dissociated a total of 80 adult fly abdominal cuticles (along with the attached adipose tissue and oenocytes) and subjected the single nuclei to 10X genomics-based (Zheng et al., 2017) single-nuclei RNA-sequencing (snRNA-seq) (Figure 2A). Two independent rounds of sequencing were performed on two biological replicates (with 40 animals per replicate) to obtain a median read depth of 8904 reads per nucleus (Figure S2A). Because tissue dissociations for the single nuclei preparations are often associated with the risk of ambient RNA contamination, our quality control pipeline included SoupX (Young and Behjati, 2018) to eliminate potential ambient RNA from our analysis. Further, we used Harmony (Korsunsky et al., 2019) that is integrated into Seurat (Stuart et al., 2019) to correct for batch effects in the two replicates to finally retain 15,280 nuclei with a median of 192 genes per nucleus for downstream analysis (Figure S2-B,D; Table S1). Our clustering analysis revealed 10 unique clusters, where three major clusters were assigned to adipose tissue, oenocytes, and muscle based on known markers for each of these tissue types including *apolpp*, *fasn3*, and *mhc*, respectively (Figure 2B; S2E; Table S2). We validated our snRNA-seq data using GAL4 lines for certain top enriched and novel genes such as *Pellino* (*Pli*), *sallimus* (*sls*), and *geko* and found that they specifically express in adipose tissue, muscle, and oenocytes, respectively (Figure S2F-I). With regards to the rest of the minor clusters (4-10), we believe most of them are remnant tissues most likely pertaining to gut/malpighian tubule based on the enrichment of *alphaTry*, *Whe*, and *Mur18B* (Clusters 4-6, respectively; Table S2). On the other hand, we consider clusters 7-10 likely to be part of the ejaculatory bulb (Eb) as they are enriched in certain male specific genes such as *bond*, *EbpIII*, *soti*, and *Ebp*, respectively (Figure S2J; Table S2).

**Figure 2:**
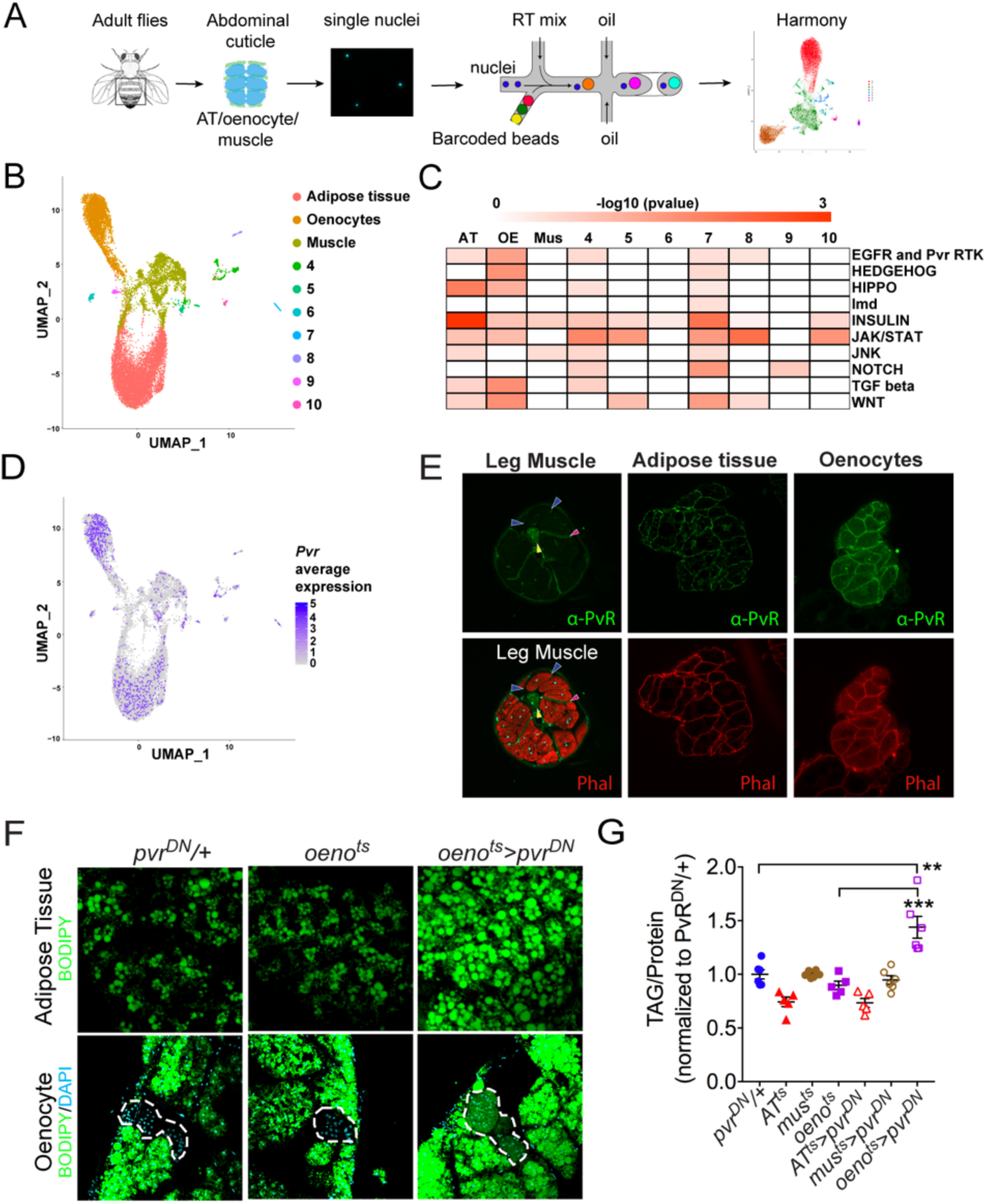
Single Nuclei-RNA-sequencing reveals that oenocyte-specific PvR signaling protects against obesity (A) Schematic of snRNA-seq workflow. Adult fly abdomens are dissected and dissociated to obtain high quality single nuclei for downstream encapsulation by 10X genomics-based snRNA-seq platform and subsequent sequencing and analysis using Harmony. (B) Uniform Manifold Approximation and Projection (UMAP) plot representing 10 unique clusters. Each color and dot in the plot represent a cluster and a single nucleus, respectively. (C) Pathway enrichment analysis reveals EGFR and Pvr Receptor Tyrosine Kinase (RTK) signaling pathway enriched in oenocytes (OE) when compared to other clusters including Adipose Tissue (AT) and Muscle (Mus). (D) UMAP plot representing the average expression of *Pvr*, which is highly enriched in oenocytes and to a lesser extent in adipose tissue and muscle. (E) Anti-PvR (Green) and phalloidin-594 (Red) staining of *w^1118^* (VDRC isogenic stock) adult male leg musculature (cross-section), adipose tissue and oenocytes. Yellow and red arrowheads show the leg axon bundle and trachea respectively. Blue arrowheads show the sarcolemma of individual leg muscle bundles. (F) BODIPY staining showing neutral lipid accumulation in adult male adipose tissue and oenocytes (dorsal abdominal cuticle) of control males and males over-expressing *pvr^DN^* in the adult oenocytes (*oeno^ts^>pvr^DN^)*. (G) Total TAG content of adult males over-expressing *pvr^DN^* in the adult adipose tissue (*AT^ts^>pvr^DN^*), muscle (*mus^ts^>pvr^DN^*) and oenocytes (*oeno^ts^>pvr^DN^)*. Flies containing the UAS construct and the tissue specific drivers alone serve as controls. n=5-6, One-Way ANOVA followed by Tukey’s HSD test.

To explore the pathways that are enriched in each of these clusters, we performed pathway enrichment analysis (Fig. 2-C). Interestingly, we found that EGFR and PVR RTK signaling pathway is highly enriched in the oenocytes (Figure 2C; Table S3). While the expression of *Egfr* is specifically enriched in the adipose tissue (Figure S2J), we identify the Pvf1 receptor Pvr to be highly enriched in the oenocytes, albeit a mild enrichment seen in the other clusters (Figure 2D). We examined the distribution of PvR in the major metabolic tissues of interest: muscle, adipose tissue and the oenocytes, by immunostaining with an anti-PvR antibody (Rosin et al., 2004). Consistent with the prediction from the snRNA-seq analysis, PvR is present most prominently on the surface of the oenocytes followed by the adipose tissue cells (Fig. 2-E). In the leg muscles and indirect flight muscles, PvR localizes to the muscle sarcolemma (Fig. 2-E and S3-B), although the level of the protein on the sarcolemma of the leg muscles is relatively weak (Figure 2E).

### Oenocyte-specific loss of Pvf-Receptor (PvR) signaling leads to obesity

Since our data shows PvR and PvR-signaling to be enriched in the oenocytes, we asked whether PvR signaling in the oenocyte is necessary for protecting the adult flies against obesity. To test this possibility, we inhibited PvR signaling specifically in the oenocyte and determined the effect on whole animal TAG levels and lipid accumulation in the adipose tissue and oenocytes. Impairing PvR signaling in the adult oenocytes, by over-expressing a dominant negative form of the receptor, *pvr^DN^*, (*oeno^ts^>pvr^DN^*) (Brückner et al., 2004) led to obesity phenotypes similar to *mus^ts^>Pvf1-i* flies (Fig. 2-F,G). Similarly, impairing PvR signaling in the oenocyte by expressing an RNAi against *pvr (oeno^ts^>pvr-i)* also leads to obesity (Fig. S3-D). The obesity phenotype was also observed in *oeno^ts^>pvr^DN^* female flies, indicating that the phenotype is not caused by loss of PvR signaling in the male accessory glands where the *PromE-Gal4* driver is also expressed (Fig. S3-C) (Billeter et al., 2009). Surprisingly, over-expressing *pvr^DN^* in the adult adipose tissue and muscle did not lead to an obesity phenotype indicating that Pvf/PvR signaling is primarily required in the oenocytes to regulate lipid abundance (Fig. S3-A). These results suggest that muscle-derived Pvf1 signals specifically to the oenocytes to regulate lipid content of the adipose tissue and steatosis in the oenocytes.

### Oenocyte-specific loss of mTOR signaling leads to obesity

Downstream of PvR, Pvf signaling primarily activates the Ras/Raf/MEK/ERK pathway. To determine whether oenocyte-specific ERK signaling regulates lipid homeostasis, we measured neutral lipid storage in *oeno^ts^>ERK-i* flies. Two independent and validated RNAi transgenes against ERK failed to replicate the obesity phenotype observed in *oeno^ts^>pvr^DN^* flies, indicating that PvR signaling in the oenocytes regulates lipid levels via an ERK-independent mechanism (Fig. 3-A,C). Previous studies in *Drosophila* S2 and Kc cells have shown that PvR can also activate the mTOR pathway (Sopko et al., 2015; Tran et al., 2013). To test whether oenocyte-specific mTOR signaling is involved in regulating lipid homeostasis, we inhibited mTOR signaling in the oenocytes by over-expressing both *tsc1* and *tsc2* (*oeno^ts^>tsc1,tsc2*). Similar to *mus^ts^>pvf1-i* and *oeno^ts^>pvr^DN^* flies, *oeno^ts^>tsc1, tsc2* flies showed massive accumulation of neutral lipids in both the adipose tissue and the oenocytes (Fig. 3-B,C). Similar to vertebrates, *Drosophila* mTOR pathway can be activated by insulin/Pi3K/Akt signaling. To determine the potential involvement of oenocyte-specific insulin signaling in regulating lipid homeostasis, we over-expressed a dominant negative form of InR (*oeno^ts^>inr^DN^*) and examined the effect on lipid accumulation. *oeno^ts^>inr^DN^* flies did not show any increase in accumulation of lipids either in the adipose tissue or the oenocytes compared to control flies (Fig. 3-D,E). We further investigated whether PvR, being a receptor tyrosine kinase, can activate mTOR signaling in the oenocyte via activation of Pi3K/Akt1 and regulate lipid homeostasis. The *Drosophila* genome encodes for three Pi3Ks (Pi3K92E, Pi3K59F and Pi3K68D) and one regulatory subunit (Pi3K21B). We knocked down each of the Pi3K components in the oenocytes and determined the effect on lipid accumulation in the oenocytes and adipose tissue (Fig. S4). Oenocyte-specific loss of either Pi3K92E and the regulatory subunit Pi3K21B led to the steatosis phenotypes (Fig. 3-D,E). Additionally, oenocyte-specific loss of *akt1* also led to the steatosis phenotypes indicating that the Pi3K-Akt1 pathway in the oenocytes regulate lipid homeostasis (Fig. 3-D,E). Taken together, these results reveal that Pi3K/Akt1/mTOR signaling in the oenocyte protects against obesity.

**Figure 3:**
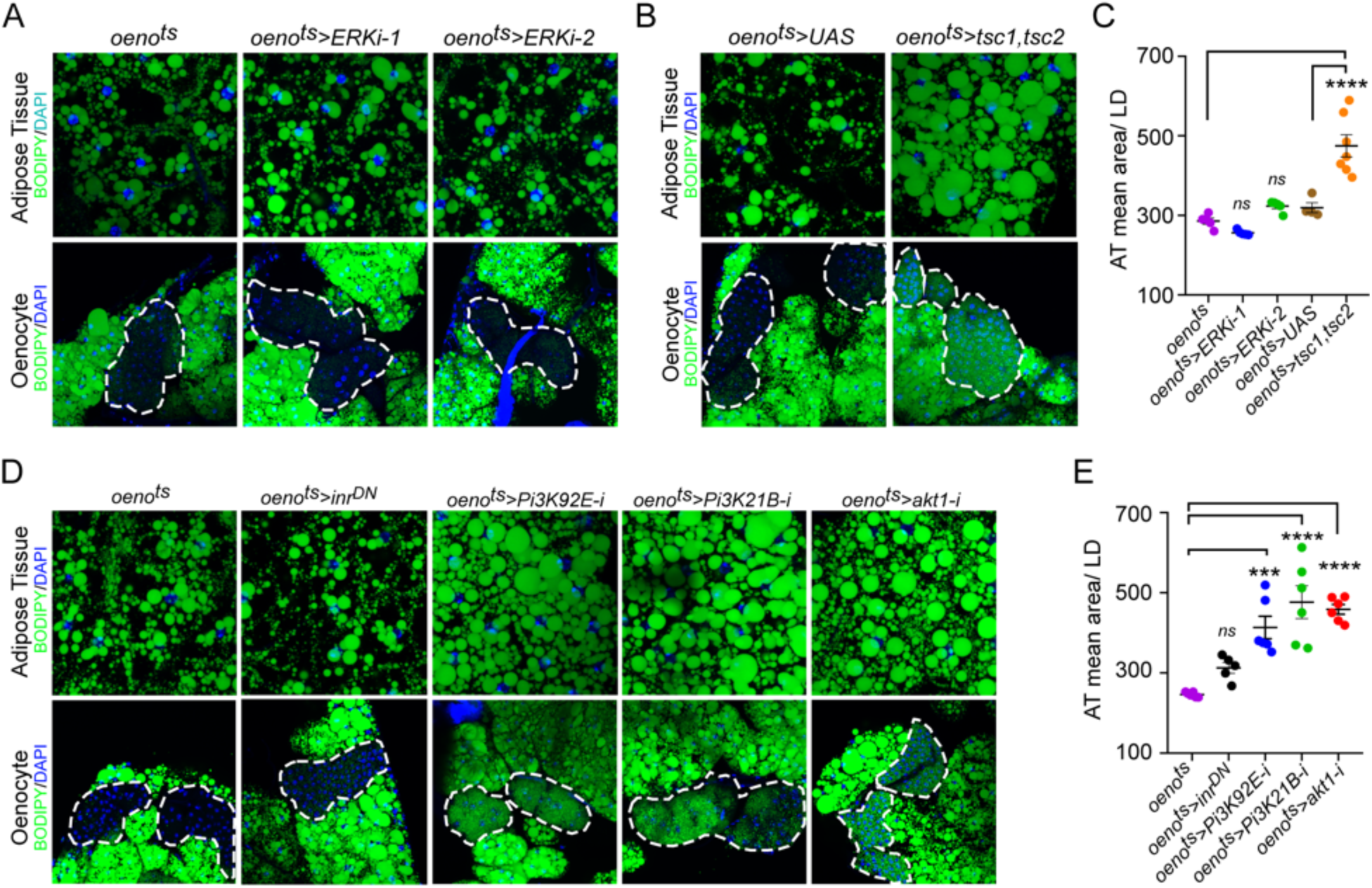
Oenocyte-specific Pi3K/Akt1/mTOR signaling protects against obesity (A) BODIPY staining showing neutral lipid accumulation in the adipose tissue and oenocytes of control males (*oeno^ts^=oenocyte-Gal4^Gal80ts^*) and males with oenocyte-specific knockdown of ERK (*oeno^ts^>ERKi-1* and *oeno^ts^>ERKi-2*) using two independent RNAi lines. (B) BODIPY staining showing neutral lipid accumulation in the adipose tissue and oenocytes of control males (*oeno^ts^*) and males with oenocyte-specific over-expression of *tsc1* and *tsc2* (*oeno^ts^>tsc1,tsc2*). (C) Mean lipid droplet size (≥10 microns in diameter) in the adipose tissue of flies shown in fig. 2A and B. Only oenocyte-specific loss of mTOR signaling (*oeno^ts^>tsc1,tsc2*) show a significant increase (*p<0.001*) compared to the control genotype. n=6, One-Way ANOVA followed by Tukey’s HSD test. (D) BODIPY staining showing neutral lipid accumulation in the adipose tissue and oenocytes of control males (*oeno^ts^*) and males with oenocyte-specific over-expression of *inr^DN^* (*oeno^ts^>inr^DN^*) and oenocyte-specific (*oeno^ts^>*) knockdown of *Pi3K92E* (*Dp110*), *Pi3K21B (Dp60)* and *akt1*. (E) Mean lipid droplet size (≥10 microns in diameter) in the adipose tissue of flies shown in fig. 2D. Oenocyte-specific (*oeno^ts^>*) knockdown of *Pi3K92E* (*Dp110*), *Pi3K21B (Dp60),* and *akt1* lead to a significant increase (*p<0.001* for *Pi3K92E,* and, *p<0.0001* for *Pi3K21B* and *akt1*) compared to the control genotype. n=6, One-Way ANOVA followed by Tukey’s HSD test.

### mTOR signaling acts downstream of PvR in the oenocytes to regulate systemic lipid stores

We next analyzed whether PvR signals through mTOR in the oenocytes to regulate lipid metabolism. For this, we first measured the levels of phospho-4EBP (p4EBP), a direct target of mTOR, in the oenocytes of *mus^ts^>pvf1-i* and *oeno^ts^>pvr^DN^* flies. Both *mus^ts^>pvf1-i* flies and *oeno^ts^>pvr^DN^* flies showed a strong and significant down-regulation of p4EBP signal in the oenocytes compared to Gal4 alone controls (Fig. 4-A,B,C), indicating that muscle Pvf regulates TORC1 signaling in the oenocyte.

**Figure 4:**
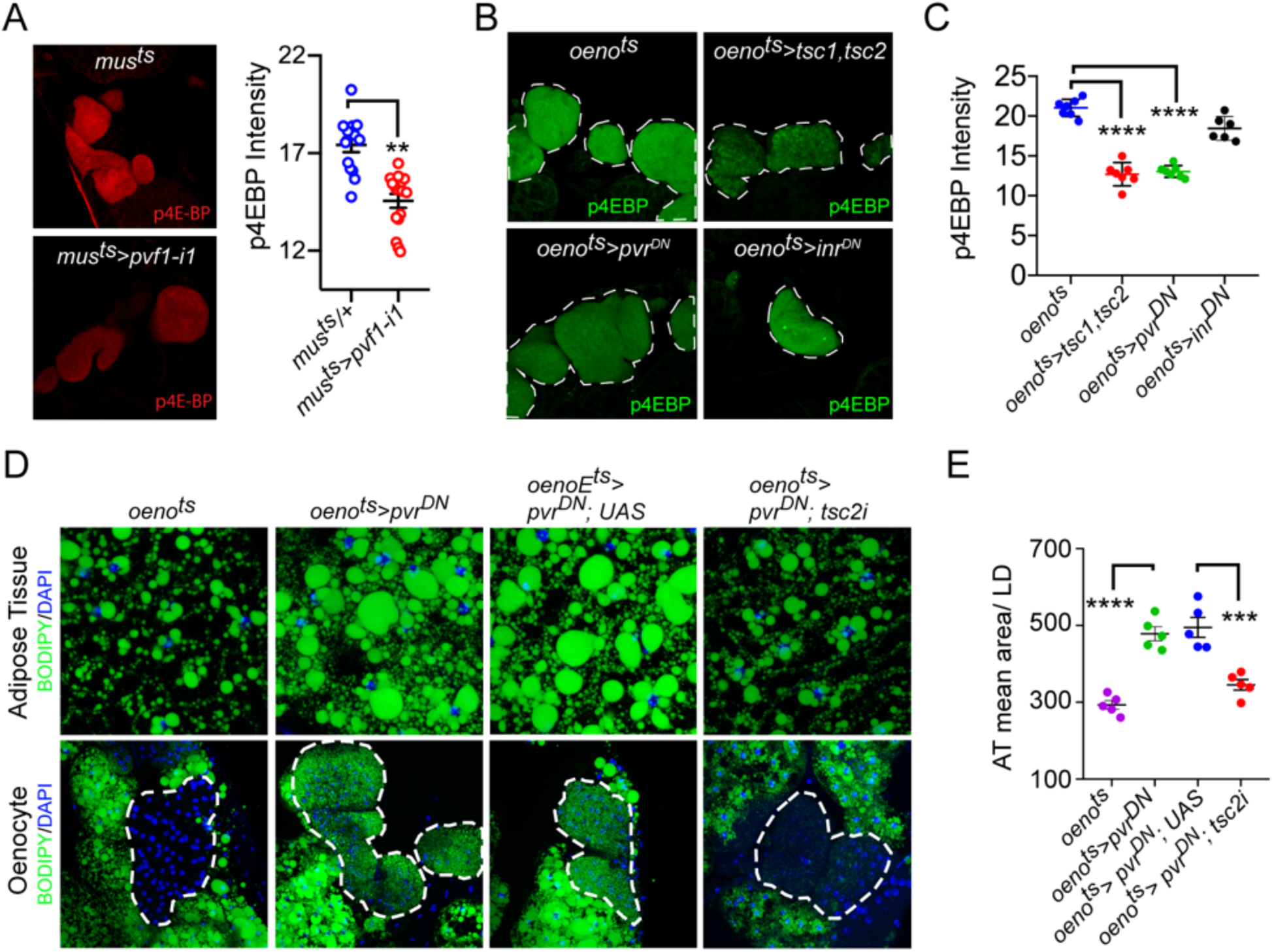
Pi3K/Akt1/mTOR signaling acts downstream of PvR in the oenocytes to regulate adiposity (A) p4EBP staining (Red) of the oenocytes from control (*mus^ts^=muscle-Gal4^Gal80ts^*) and flies with muscle-specific knockdown of *pvf1* (*mus^ts^=pvf1-i1*). Muscle-specific loss of *pvf1* leads to a significant decrease (*p<0.01*) in p4EBP levels in the oenocytes. n=10, student t-test (B) p4EBP staining (Green) of oenocytes from control flies (*oeno^ts^=oenocyte-Gal4^Gal80ts^*) and flies with oenocyte-specific over-expressing *tsc1/tsc2* (*oeno^ts^>tsc1,tsc2*), *pvr^DN^* (*oeno^ts^*>*pvr^DN^*) or *inr^DN^* (*oeno^ts^>inr^DN^*). (C) Quantification of p4EBP staining intensity in the oenocytes for samples shown in fig. 3C. Oenocyte-specific over-expression of *tsc1/tsc2* (*oeno^ts^>tsc1,tsc2*) and *pvr^DN^* (*oeno^ts^*>*pvr^DN^*) led to a significant reduction in p4EBP levels (*p<0.0001*). Over-expression of *inr^DN^* (*oeno^ts^>inr^DN^*) however does not affect p4EBP levels significantly. n = 6/7, Student t-test (D) BODIPY staining showing neutral lipid accumulation in the adipose tissue and oenocytes of control males (*oeno^ts^*) and males with oenocyte-specific over-expression of either *pvr^DN^* (*oeno^ts^*>*pvr^DN^*) or over-expression of *pvr^DN^* along with *tsc2* knockdown (*oeno^ts^*>*pvr^DN^,tsc2-i*). Flies over-expressing *pvr^DN^* in the oenocytes along with an empty UAS construct (*oeno^ts^*>*pvr^DN^,UAS*) serve as an additional control to account for any effect of Gal4 dilution on the obesity phenotype. (E) Mean lipid droplet size (≥10 microns in diameter) in the adipose tissue of flies shown in fig. 3A. Oenocyte-specific knockdown of *tsc2* along with *pvr^DN^* over-expression (*oeno^ts^*>*pvr^DN^,tsc2-i*) significantly rescues the obesity phenotype observed flies with oenocyte-specific over-expression of *pvr^DN^* (*p<0.001*). n=5, One-Way ANOVA followed by Tukey’s HSD test.

Consistently, the extent of p4EBP down-regulation is similar to what is observed in *oeno^ts^>tsc1, tsc2* flies that were used as a positive control for the assay (Fig. 4-B). Interestingly, oenocyte-specific loss of insulin receptor signaling (*oeno^ts^>inr^DN^*) failed to affect p4EBP levels, further supporting the observation that PvR, but not InR, activates mTOR signaling in the oenocytes of well-fed flies (Fig. 4-B). In addition, we tested whether the obesity phenotype of *oeno^ts^>pvr^DN^* flies could be rescued by activating mTOR signaling. To do this, we co-expressed *pvr^DN^* and a *tsc2-RNAi* transgene in the oenocytes (*oeno^ts^>pvr^DN^; tsc2-i*) and compared the lipid content of these flies to control fly lines. *tsc2* knockdown strongly suppressed the obesity phenotype induced by *pvr^DN^* (Fig. 4-D,E) indicating that mTOR signaling functions downstream of PvR to regulate the steatosis phenotype. Altogether, we demonstrate that in *Drosophila,* muscle-derived Pvf1 signals through PvR in the oenocyte to activate mTOR, which in turn protects the animal against obesity.

### Muscle-to-Oenocyte Pvf1 signaling regulates lipid synthesis

*Drosophila* oenocytes are known to facilitate starvation-induced lipid mobilization in the *Drosophila* larvae and loss of this tissue leads to increased starvation sensitivity (Gutierrez et al., 2007). Similarly, in adult flies the oenocytes play a role in imparting starvation resistance by regulating production of very long chain fatty acids (VLCFAs) for waterproofing of the cuticle, especially when flies are starved under lower humidity conditions, and possibly by regulating lipid mobilization (Chatterjee et al., 2014; Storelli et al., 2019). We first investigated whether muscle-to-oenocyte Pvf1 signaling plays a role in starvation resistance. Compared to control flies, *mus^ts^>pvf1-i* and *oeno^ts^>pvr^DN^* animals showed increased starvation resistance, suggesting that they are capable of mobilizing stored nutrients in response to starvation and are not defective in production of VLCFAs needed for waterproofing of the cuticle (Fig. 5-A). The improved starvation resistance of *mus^ts^>pvf1-i* and *oeno^ts^>pvr^DN^* flies most likely reflects the fact that these animals had higher levels of stored TAGs and hence were able to use these reserves for a longer duration. As starvation can induce strong catabolic signals that can easily mask minor defects in lipid mobilization in *mus^ts^>pvf1-i* and *oeno^ts^>pvr^DN^* flies, we measured the rate of lipid mobilization under steady state feeding conditions using radioisotope chasing. We labeled the TAG stores of control and experimental flies with [1-^14^C]-Oleate for 3 days.

**Figure 5:**
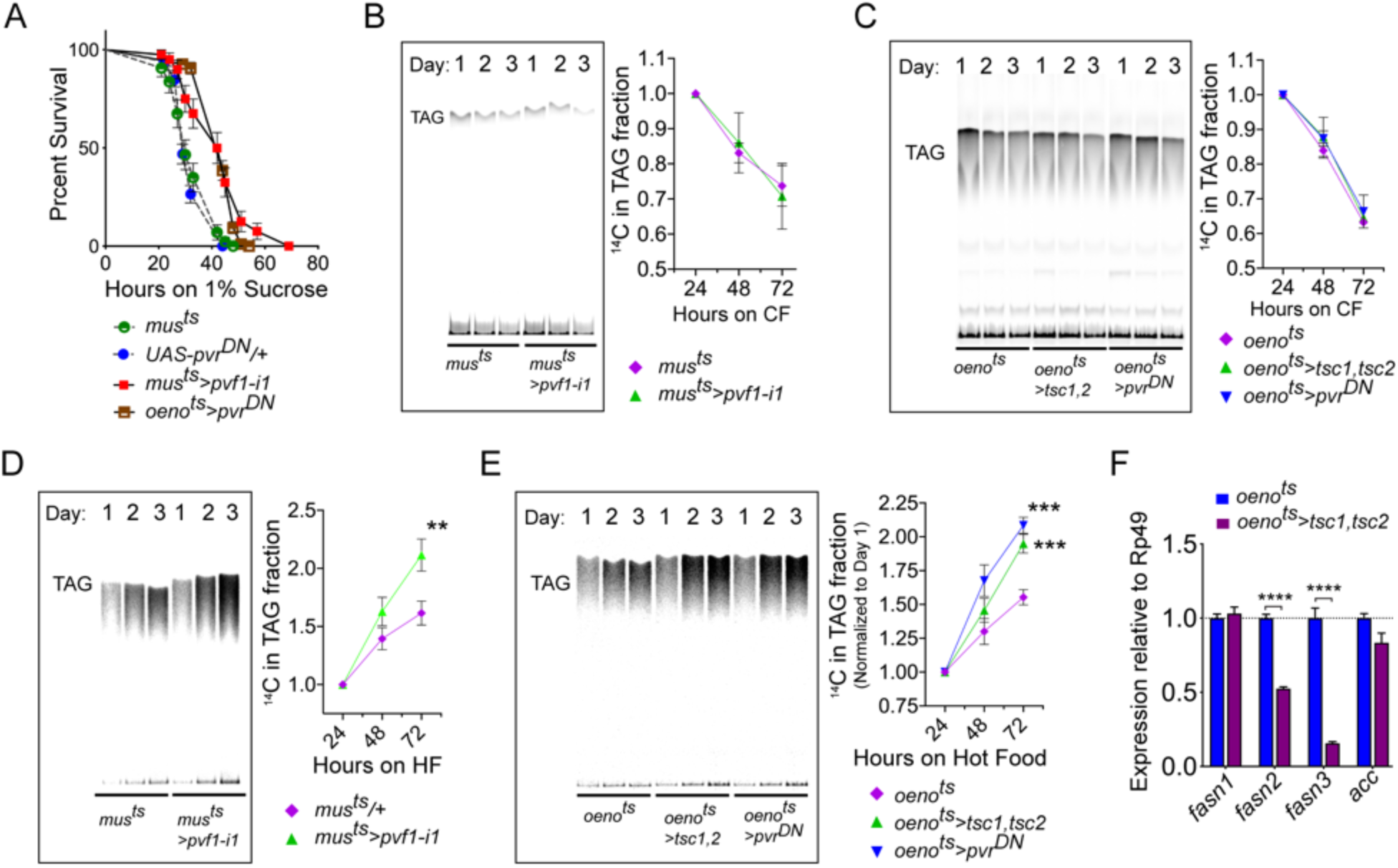
Muscle to oenocyte Pvf1 signaling suppresses lipid synthesis (A) Starvation resistance of adult male flies on 1% sucrose and 0.8% agar food. Males with muscle-specific knockdown of *pvf1* (*mus^ts^>pvf1-i1*) and oenocyte-specific loss of PvR signaling (*oeno^ts^*>*pvr^DN^*) were significantly more resistant to starvation compared to controls flies (*mus^ts^= muscle-Gal4^Gal80ts^* and *UAS-pvr^DN^/+*). n=100, Kaplan Myer log rank test. (B) Rate of lipid mobilization in control (*mus^ts^*) and flies with muscle-specific loss of *pvf1* (*mus^ts^>pvf1-i1*). n=4, Multiple Student t-test. (C) Rate of lipid mobilization in control (*oeno^ts^=oenocyte-Gal4^Gal80ts^*) and flies with oenocyte-specific loss of mTOR signaling (*oeno^ts^>tsc1,tsc2*) and PvR signaling (*oeno^ts^>pvr^DN^*). n=4, Multiple Student t-test. (D) Rate of lipid synthesis and incorporation from [U-^14^C]-Sucrose in control (*mus*^ts^) and flies with muscle-specific loss of *pvf1* (*mus^ts^>pvf1-i1*). Flies lacking Pvf1 in the muscle show a significantly faster rate of lipid incorporation compared to control animals. n=4, *p≤0.01* at 72 hours on hot food. Multiple student t-test. (E) Rate of lipid synthesis and incorporation from [U-^14^C]-Sucrose in control flies (*oeno^ts^*) and flies with oenocyte-specific loss of mTOR signaling (*oeno^ts^>tsc1,tsc2*) and PvR signaling (*oeno^ts^>pvr^DN^*). Flies lacking mTOR or PvR signaling in the oenocytes show a significantly faster rate of lipid incorporation compared to control animals. n=4, *p≤0.001* at 72 hours on hot food. Multiple student t-test. (F) Expression level of key lipid synthesis genes in control (*oeno^ts^*) and flies with oenocyte-specific loss of mTOR signaling (*oeno^ts^>tsc1,tsc2*). Only oenocyte-specific fatty acid synthases, *fasn2* and *fasn3*, show a significant reduction in expression in the experimental flies. n=4, One-Way Anova followed by Tukey’s HSD test.

Subsequently, we shifted the labeled flies to cold food and collected samples at 24, 48 and 72 hours post transfer and measured ^14^C label in the TAG fractions using thin-layer chromatography (TLC).

Interestingly, *mus^ts^>pvf1-i* flies showed similar rates of lipid mobilization from TAG stores compared to control flies (Fig. 5-B). Similarly, *oeno^ts^>tsc1, tsc2* and *oeno^ts^>pvr^DN^* flies also showed comparable rates of lipid mobilization compared to control animals, indicating that loss of the muscle-to-oenocyte Pvf1 signaling axis does not impair lipid mobilization (Fig 5-C). Since loss of the muscle-to-oenocyte Pvf1 signaling axis did not affect lipid mobilization, we tested whether flies lacking this pathway show increased lipid synthesis. To test this possibility, we transferred experimental and control animals to ^14^C-U-Sucrose containing food and measured the levels of ^14^C-incorporation over time in the TAG fraction of the flies using TLC. We found that *mus^ts^>pvf1-i*, *oeno^ts^>tsc1, tsc2,* and *oeno^ts^>pvr^DN^* flies all showed an increased rate of ^14^C incorporation into TAG fractions compared to control animals (Fig. 5-D,E). In absence of any effects on lipid mobilization, the increased rate of lipid incorporation indicates an increased rate of lipid synthesis in the experimental flies.

Since loss of mTOR signaling in the oenocytes led to increased lipid synthesis, we investigated the role of mTOR signaling in regulating lipid synthetic genes in the oenocytes. We extracted total RNA from adult oenocyte/adipose tissue complexes and measured the expression levels of two oenocyte specific fatty acid synthases, *fasn2* and *fasn3*. In addition, we measured the expression of two lipogenic genes, *acc* and *fasn1*, that are not exclusively expressed in the adult oenocytes in *Drosophila*. While expression levels of *acc* and *fasn1* did not change in *oeno^ts^>tsc1,tsc2* flies, expression levels of *fasn2* and *fasn3* were strongly downregulated (Fig. 5-F). This indicates that loss of mTOR signaling downregulates lipogenic genes in the oenocytes, and that the increased lipid synthesis observed in *oeno^ts^>tsc1,tsc2* flies is caused by a mechanism independent of the role of mTOR in regulating the expression of lipogenic genes in the oenocytes.

### Muscle-to-Oenocyte Pvf1 signaling regulates post-eclosion restoration of stored lipids in adult adipose tissue

When adult flies emerge from their pupal cases the adult adipose tissue has very low stored lipid content (Storelli et al., 2019), and adipose tissue cells of post-eclosion flies are also notably small in size (Fig. S5). Over the course of the next 3 to 7 days the adipose tissue recovers its lipid stores through feeding and *de-novo* lipid synthesis, and expands significantly in size both at cellular and tissue levels (Fig. S5). While the average size of the lipid droplets does not change drastically during this period, the number of lipid droplets per cell increases drastically and a large number of smaller lipid droplets start appearing in the adipose tissue cells (Fig. S5). These observations suggest that restoration of adipose tissue lipid stores happens by formation of new lipid droplets that become bigger in size as adipose tissue lipid recovery progresses. Since muscle-to-oenocyte Pvf1 signaling axis negatively regulates lipid synthesis, we hypothesized that this pathway is needed to inhibit lipid synthesis once the restoration of adipose tissue lipid stores reaches completion. Consistent with this hypothesis, we find that muscle-specific expression levels of *pvf1* is low in newly eclosed flies and increases rapidly over the course of the next 7 days (Fig. 6-A). To further test the hypothesis that muscle-Pvf1 limits the extent of TAG recovery in the adipose tissue of newly eclosed flies, we over-expressed *pvf1* in the adult muscle from day 1 of eclosion and measured the frequency of large lipid droplets (LD ≥5μm in diameter/cell) per cell using BODIPY staining. Compared to controls, *mus^ts^>pvf1* flies tend to accumulate much lower number of large lipid droplets per adipose tissue cell (Fig. 6-B,D). Additionally, the experimental animals tend to have larger number of empty adipose tissue cells per animal compared to control flies (Fig. 6-C).

**Figure 6:**
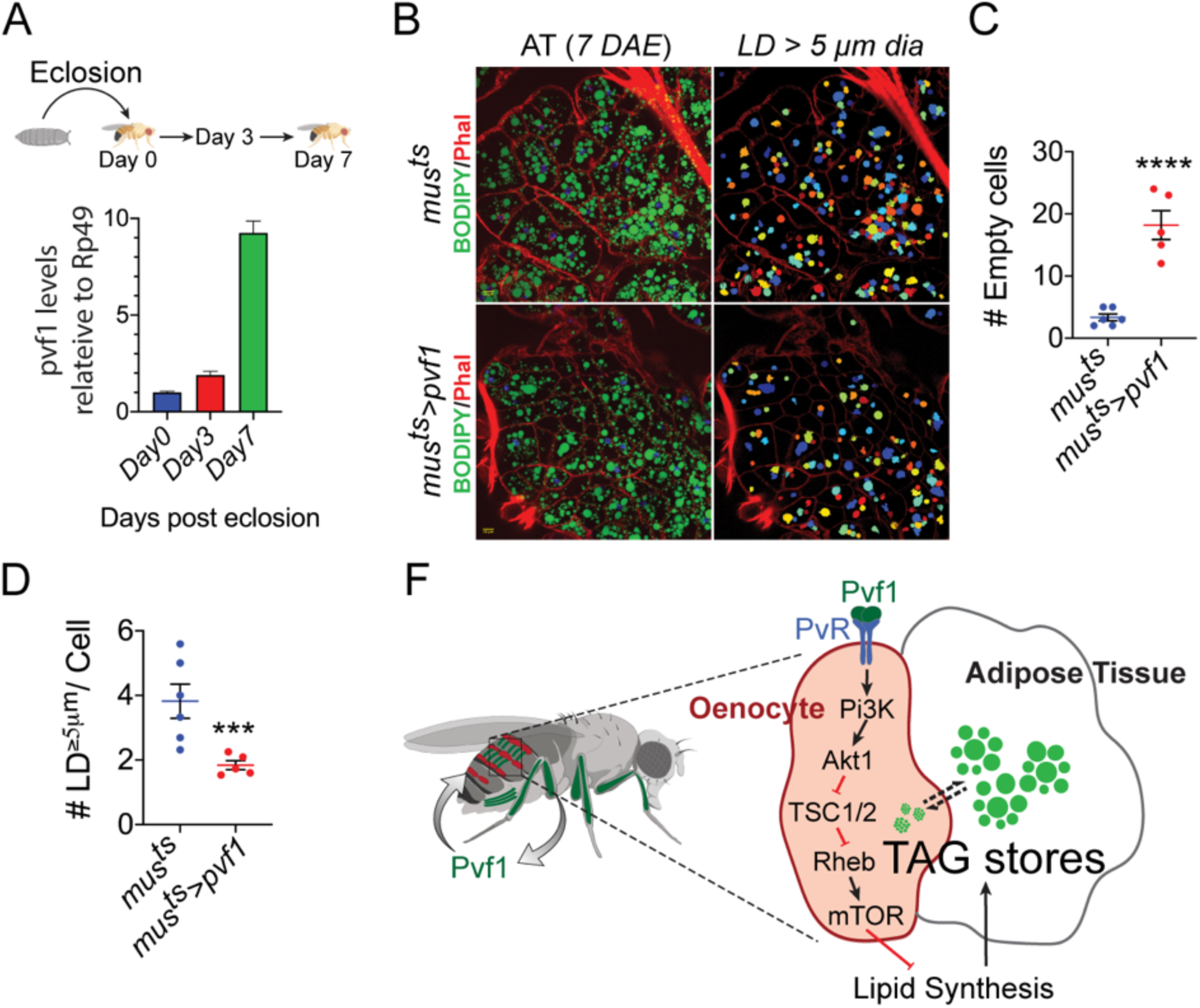
Muscle-Pvf1 limits post eclosion lipid recovery by suppressing lipid synthesis (A) Muscle-specific expression of *pvf1* in freshly eclosed *w^1118^* males on day 0, 3 and 7 post eclosion. (B) BODIPY staining of adipose tissue from 7-day old control males (*mus^ts^= muscle-Gal4^Gal80ts^*) and males with muscle-specific over-expression of *pvf1* (*mus^ts^>Pvf1*) from Day 0 of eclosion. Panels on the right show identification of lipid droplets (LD) that are ≥5 μm in diameter using Cell Profiler. (C) Quantification of number of adipose tissue cells that do not contain LDs that are ≥5 micron in diameter for control (*mus^ts^*) flies and flies with muscle-specific over-expression of *pvf1* (*mus^ts^>pvf1*). n=5/6, *p≤0.0001*, Student t-test. (D) Quantification of number of large (≥5 μm in diameter) LDs per cell in control (*mus^ts^*) flies and flies with muscle-specific over-expression of *pvf1* (*mus^ts^>pvf1*). n=5/6, *p≤0.001*, Student t-test (E) Model of the role of muscle-derived Pvf1 in regulating systemic lipid content by signaling to the oenocytes of adult male flies.

These results suggest that muscle-derived Pvf1 helps terminate the adipose tissue lipid recovery phase by suppressing lipid synthesis by signaling to the *Drosophila* oenocytes.

## Discussion

The presence in vertebrates of multiple PDGF/VEGF signaling ligands and cognate receptors makes it difficult to assess their roles in inter-organ communication. Additionally, understanding the tissue-specific roles of these molecules, while circumventing the critical role they play in regulating tissue vascularization, is equally challenging in vertebrate models. Here, we investigated the tissue specific roles of the ancestral PDGF/VEGF-like factors and the single PDGF/VEGF-receptor in *Drosophila* in lipid homeostasis. Our results demonstrate that in adult flies the PDGF/VEGF like factor, Pvf1, is a muscle-derived signaling molecule (myokine) that suppresses systemic lipid synthesis by signaling to the *Drosophila* hepatocyte-like cells/oenocytes. Newly eclosed adult flies emerge from their pupal case with limited lipid stores in their adipose tissue. While the larval adipose tissue cells persists through metamorphosis, much of the lipids released from these cells are used up to form the water resistant barrier on their exoskeleton that protects them from dehydration (Storelli et al., 2019). The adipose tissue lipid stores are restored after emergence through feeding and de-novo lipid synthesis. This rate of lipid incorporation is quite high during the first 3-7 days of the adult life. However, at the end of this adipose tissue lipid restoration phase, the rate of lipid synthesis must be suppressed to avoid over-loading of the adipose tissue and prevent the consequent effects of lipid toxicity. Our data indicates that muscle Pvf1 signaling suppresses lipid synthesis at the end of the lipid recovery phase by activating mTOR signaling in the oenocytes.

### *Drosophila* Pvf1 functions as a myokine that suppresses lipid synthesis

Our study reveals that Pvf1 is abundant in the tubular muscles of the *Drosophila* leg and abdomen. In these striated muscles the protein localizes between individual myofibrils and is particularly enriched at the M and Z bands. *Drosophila* musculature can be broadly categorized into two groups, the fibrillar muscles and the tubular muscles, with distinct morphological and physiological characteristics. *Drosophila* IFMs of the thorax belong to the fibrillar muscle group and are stretch-activated, oxidative, slow twitch muscles that are similar to vertebrate cardiac muscles (Schönbauer et al., 2011). By contrast, *Drosophila* leg muscles and abdominal muscles belong to the tubular muscle group. These muscles are striated, Ca^2+^ activated and glycolytic in nature. The tubular muscles are structurally and functionally closer to vertebrate skeletal muscles (Schnorrer et al., 2010; Schönbauer et al., 2011). Expression of Pvf1 in the tubular muscles of the *Drosophila* leg may reflect a potentially conserved role of this molecule as a skeletal-muscle derived myokine. The fact that most of the myokines in vertebrates were identified in striated skeletal muscles supports this possibility (Pedersen and Febbraio, 2012; So et al., 2014). Moreover, vertebrate VEGF ligands, VEGF-A and VEGF-B, have also been shown to be stored and released from skeletal muscles (Boström et al., 2012; Vind et al., 2011).

Interestingly, in vertebrates, the expression and release of VEGF ligands are regulated by muscle activity (Boström et al., 2012; Vind et al., 2011). In mice, expression of VEGF-B in the skeletal muscles is regulated by PGC1-α, one of the key downstream effectors of muscle activity. Additionally, expression of VEGF-B is upregulated in both mouse and human skeletal muscles in response to muscle activity (Boström et al., 2012; Vind et al., 2011). Similarly, expression of VEGF-A is induced by muscle contraction (Boström et al., 2012). We did not see any effect of muscle activity on the expression levels of *pvf1* in the *Drosophila* muscles. We also could not demonstrate whether muscle activity regulates release of Pvf1 primarily due to the difficulty in collecting adequate amounts of hemolymph from the adult males. However, the localization of Pvf1 to the M/Z bands suggests a potential role for muscle activity in Pvf1 release. The M and Z bands of skeletal muscles are important centers for sensing muscle stress and strain. These protein-dense regions of the muscle house a number of proteins that can act as mechano-sensors and mediate signaling events including translocation of selected transcriptional factors to the nucleus (Hoshijima, 2006; Lange et al., 2019; Miller et al., 2003). Pvf1, therefore, is ideally located to be able to sense muscle contraction and be released in response to muscle activity. Further work, contingent on the development of new tools and techniques, will be necessary to measure Pvf1 release into the hemolymph and study the regulation of this release by exercise.

We have previously shown that Pvf1 released from gut tumors generated by activation of the oncogene *yorkie* leads to wasting of *Drosophila* muscle and adipose tissue (Song et al., 2019). Adipose tissue wasting in these animals is characterized by increased lipolysis and release of free fatty acids (FFAs) in circulation. However, we did not observe any role of Pvf signaling in regulating lipolysis in the adipose tissue of healthy well-fed flies without tumors. Loss of PvR signaling in the adipose tissue did not have any effect on lipid content. Additionally, over-expressing Pvf1 in the muscle did not lead to the bloating phenotype commonly seen in cachectic animals with gut tumors (Kwon et al., 2015; Song et al., 2019). We conclude that Pvf1 affects wasting of the adipose tissue only in the context of gut tumors and that the effect could involve the following mechanisms: 1) the gut tumor releases pathologically very high levels of Pvf1 into circulation leading to ectopic activation of PvR signaling in the adipose tissue, and, that such high levels of Pvf1 are not released by the muscle (even when *pvf1* is over-expressed in the muscle); and 2) Pvf1 causes adipose tissue wasting in the context of other signals that emanate from the gut tumor that are not available in flies that do not have tumors.

### *Drosophila* oenocytes regulate lipid synthesis and lipid content of the adipose tissue

Only oenocyte-specific loss of PvR signaling phenocopies the obesity phenotype caused by muscle-specific loss of Pvf1, indicating that muscle-Pvf1 primarily signals to the oenocytes to regulate systemic lipid content. Additionally, muscle-specific loss of Pvf1, as well as oenocyte-specific loss of PvR and its downstream effector mTOR, leads to an increase in the rate of lipid synthesis. These observations indicate a role for the *Drosophila* oenocytes in lipid synthesis and lipid accumulation in the adipose tissue. Oenocytes have been implicated in lipid metabolism previously and these cells are known to express a diverse set of lipid metabolizing genes including but not limited to fatty acid synthases, fatty acid desaturases, fatty acid elongases, fatty acid β-oxydation enzymes and lipophorin receptors (reviewed in (Makki et al., 2014)). Functionally, the oenocytes are proposed to mediate a number of lipid metabolism processes. Oenocytes tend to accumulate lipids during starvation (presumably for the purpose of processing lipids for transport to other organs and generation of ketone bodies) and are necessary for starvation induced mobilization of lipids from the adipose tissue (Gutierrez et al., 2007; Makki et al., 2014). This role is similar to the function of the liver in clearing FFAs from circulation during starvation for the purpose of ketone body generation and redistribution of FFAs to other organs by converting them to TAG and packaging into very-low density lipoproteins (Nguyen et al., 2008). However, our [1-^14^C]-oleate chase assay did not show any effect of oenocyte-specific loss of PvR/mTOR signaling on the rate of lipid utilization, indicating that this pathway does not affect oenocyte-dependent lipid mobilization.

Oenocytes also play a crucial role in the production of VLCFAs needed for waterproofing of the cuticle (Storelli et al., 2019). Results of our starvation resistance assay indicate that loss of the muscle-to-oenocyte Pvf1 signaling axis does not affect waterproofing of the adult cuticle. Storelli et al. have recently shown that the lethality observed in traditionally used starvation assays is largely caused by desiccation unless the assay is performed under saturated humidity conditions. Since our starvation assay was performed under 60% relative humidity (i.e. non-saturated levels), it is likely that desiccation played a partial role in causing starvation-induced lethality. Any defects in waterproofing of the adult cuticle would have led to reduced starvation resistance. However, both muscle-specific loss of Pvf1 and oenocyte-specific loss of PvR led to increased starvation resistance suggesting normal waterproofing in these animals. The increased starvation resistance in these animals is likely the result of these animals having higher stored lipid content that helps them to survive longer without food.

Insect oenocytes were originally believed to be lipid synthesizing cells because they contain wax-like granules (Makki et al., 2014). These cells express a large number of lipid-synthesizing genes and the abundance of smooth endoplasmic reticulum further suggest a role for this organ in lipid synthesis and transport (Chatterjee et al., 2014; Jackson and Locke, 1989; Wigglesworth, 1988). However, evidence for potential involvement of the oenocytes in regulating lipid synthesis and lipid deposition in the adipose tissue has been lacking. The fact that two of the three fatty acid synthases (*fasn2* and *fasn3*) encoded by the *Drosophila* genome are expressed specifically in adult oenocytes suggests a potential role for these cells in lipid synthesis (Chung et al., 2014). Our observation that oenocyte-specific loss of PvR and its downstream effector mTOR leads to increased lipid synthesis and increased lipid content of the adipose tissue strongly supports this possibility. Our data further suggests that involvement of the oenocytes in mediating lipid synthesis is more pronounced in newly eclosed adults when the adipose tissue needs to actively replenish its lipid stores. In later stages of life, the lipid synthetic role of the oenocytes is repressed by the muscle-to-oenocyte Pvf1 signaling axis. Our observation also raises the question of whether FFAs made in the oenocytes can be transported to the adipose tissue for storage. We tested this possibility by over-expressing the lipogenic genes *fasn1* and *fasn3*, which regulate the rate limiting steps of de-novo lipid synthesis, in the oenocytes. We found that excess lipids made in the oenocytes do end up in the adipose tissue of the animal leading to increased lipid droplet size in the adipose tissue (Fig S6). Taken together, these results provide evidence for the role of *Drosophila* oenocytes in lipid synthesis and storage of neutral lipids in the adipose tissue of the animal. Interestingly, the vertebrate liver is also one of the primary sites for de-novo lipid synthesis and lipids synthesized in the liver can be transported to the adipose tissue for the purpose of storage (Gibbons et al., 2000; Meex and Watt, 2017). Hence, the fundamental role of the oenocytes and the mammalian liver converge with respect to their involvement in lipid synthesis.

### A unique role of oenocyte-specific mTOR signaling in lipid synthesis

We observed that oenocyte-specific loss of the components of the Pi3K/Akt1/mTOR signaling pathway leads to increased lipid synthesis. Moreover, the *Drosophila* insulin receptor did not play any role in activating mTOR signaling in the oenocytes. The increased rate of lipid synthesis in flies lacking mTOR signaling in the oenocytes is paradoxical to our current knowledge of how mTOR signaling affects expression of lipid synthesis genes. In both vertebrates and flies mTOR signaling is known to facilitate lipid synthesis by inducing the expression of key lipid synthesis genes such as *acetyl CoA-carboxylase* and *fatty acid synthase* via activation of SREBP-1 proteins (Han and Wang, 2018; Heier and Kühnlein, 2018; Porstmann et al., 2008). Interestingly, insulin dependent activation of SREBP-1 by mTOR is not universal. For instance, in the specialized cells of non-obese mouse liver, the InR does not play any role in activation of mTOR and downstream activation of SREBP-1c (Haas et al., 2012). We therefore checked how oenocyte-specific loss of mTOR signaling affects expression of oenocyte-specific fatty acid synthases (*fasn2* and *fasn3*) and oenocyte non-specific fatty acid synthetic genes (*fasn1* and *acc*). We observed that oenocyte-specific loss of mTOR strongly down-regulates only *fasn2* and *fasn3,* showing that mTOR signaling indeed positively regulates expression of lipogenic genes in the oenocytes and that mTOR is active in the oenocytes even though InR does not play a role in activating it. Rather, our data suggests that in wildtype well-fed flies mTOR signaling in the oenocytes is activated by the Pvf receptor. An increase in lipid synthesis in response to loss of mTOR in the oenocytes is quite intriguing and the mechanism remains to be addressed. This could happen either as a result of compensatory upregulation of lipid synthesis in the adipose tissue or due to disruption of an as yet unknown role of the oenocytes in lipid synthesis that hinges on mTOR signaling. The fact that the expression levels of *fasn1* and *acc* does not change significantly in animals lacking mTOR signaling in the oenocytes indicates that compensatory upregulation of lipid synthesis, if present, does not happen through transcriptional upregulation of lipid synthesis genes in the adipose tissue. It is still possible, however, that the increase in lipid synthesis is caused by post-translational modifications of the enzymes. Understanding the tissue specific alterations in gene expression and changes in the phosphorylation states of key lipogenic proteins in the adipose tissue of animals lacking mTOR signaling in the oenocytes will be necessary to parse out the mechanism.

Serum levels of VEGF-A is high in obese individuals and drops rapidly in response to bariatric surgery, suggesting a role for VEGF-A in obesity (García de la Torre et al., 2008; Loebig et al., 2010; Silha et al., 2005). However, evidence on whether VEGF-A or other VEGFs are deleterious vs beneficial in the context of the pathophysiology of obesity is unclear. Adipose tissue specific over-expression of both VEGF-B and VEGF-A has been shown to improve adipose tissue vascularization, reduce hypoxia, induce browning of fat, increase thermogenesis and protect against obesity (Elias et al., 2012; Robciuc et al., 2016; Sun et al., 2012; Sung et al., 2013). At the same time, blocking VEGF-A signaling in the adipose tissue of genetically obese mice leads to reduction of body weight gain, improvement in insulin sensitivity and a decrease in adipose tissue inflammation (Sun et al., 2012).

Moreover, systemic inhibition of VEGF-A or VEGF-B signaling by injecting neutralizing monoclonal antibodies have also shown remarkable effects in improving insulin sensitivity in the muscle, adipose tissue and the liver of high-fat diet-induced mouse models of obesity and diabetes (Hagberg et al., 2012; L. E. Wu et al., 2014). Although the evidence on the roles of VEGF/PDGF signaling ligands in obesity and insulin resistance is well established, the mechanisms clearly are quite complex and are often context dependent. Consequently, a wider look at various tissue specific roles of PDGF/VEGF signaling will be necessary to comprehensively understand the roles of PDGF/VEGF signaling in lipid metabolism. Our work demonstrates an evolutionarily conserved role for PDGF/VEGF signaling in lipid metabolism and a non-endothelial cell dependent role of the signaling pathway in lipid synthesis. Additionally, our findings suggest an atypical tissue-specific role of mTOR signaling in suppressing lipid synthesis at the level of the whole organism. Further studies will be required to determine whether vertebrate VEGF/PDGF and mTOR signaling exerts similar roles either in the vertebrate liver or in other specialized organ.

### A transcriptomic resource of adipose/oenocyte/muscle tissues

We made use of snRNA-Seq technology to identify expression of Pvr precisely in certain tissues in the complex abdominal region, which harbors several metabolically active tissues including adipose tissues, oenocytes, and muscle in *Drosophila*. As yet, there is no systematic study of a complete transcriptomics resource of each of these tissues considering the difficulty in dissecting and segregating these tissues for downstream sequencing. Thus, our study also provides a rich resource of gene expression profiles, paving way for a systems-level understanding of each of these tissues in *Drosophila*.

## Acknowledgements

The authors would like to thank the Bloomington Drosophila Stock Center (BDSC), Vienna Drosophila Resource Center (VDRC), Fly stocks of National Institute of Genetics (NIG-Fly) and the Transgenic RNAi project (TRiP) for fly stocks; the Microscopy Resources on the North Quad (MicRoN) for access to their laser scanning confocal microscopes; the Biorender team (https://biorender.com) for help with making illustrations for Figure 6F; Dr. Chandramohan Chitraju for his expert advice on designing and running the TLC experiments; Dr. Tobias C. Walther and Dr. Robert V. Farese for allowing us to use their Typhoon scanner and Dr. Stephanie S. Mohr, Dr. Patrick Jouandin and Dr. Ben Ewen-Campen for their comments on the manuscript. V.B. and S.H.S. were supported by funds from the Harvard Medical School Tools and Technology Committee. A.C.G. was supported by a postdoctoral fellowship from the American Heart Association (18POST33990414). This work was supported by the National Institute of Health (5RO1AR05735210 and P01CA120964). N.P. is an investigator of the Howard Hughes Medical Institute.

## Author contributions

A.C.G. and S.G.T. performed experiments; A.C.G., S.G.T., Y.L., Y.H., S.H.S., V.B. and N.P. analyzed and discussed the data; A.C. and Y.H. developed the web portal, A.C.G., S.G.T. and N.P. wrote the manuscript. A.C.G. and N.P conceived and supervised the project.

## Materials and Methods

### Drosophila strains

A detailed list of fly strains and genotypes used for each figure is provided in the Key Resources Table. For tissue specific transgene expression, we used the following temperature sensitive strains. *PromE-Gal4, tub-Gal80^ts^* (BDSC:65406) for the oenocytes (*oeno^ts^*), *tub-Gal80^ts^; mef2-Gal4* and *tub-Gal80^ts^; mhc-Gal4* for the muscle (*mus^ts^*), *tub-Gal80^ts^; Lpp-Gal4* for the adipose tissue (*AT^ts^*)and *myo1A-Gal4; tub-Gal80^ts^* for the gut (*gut^ts^*). Control animals throughout the paper were generated by either crossing the temperature sensitive driver lines to the isogenized *w^1118^* flies from VDRC or by crossing relevant UAS-lines to the isogenized *w^1118^* flies from VDRC (VDRC60000). The *UAS-fasn3* line was generated in the lab. For gene silencing we used RNAi lines from the TRiP (https://fgr.hms.harvard.edu/fly-in-vivo-rnai) available through BDSC, NIG-Fly (https://shigen.nig.ac.jp/fly/nigfly/) and VDRC (https://stockcenter.vdrc.at/control/main.

### Fly food and temperature

Flies strains were routinely kept at 25 °C or 18°C on standard lab food (SF) composed of 15g yeast, 8.6g soy flour, 63g corn flour, 5g agar, 5g malt, 74 ml corn syrup per liter. 15 w/v HSD was prepared by adding 10g of Sucrose to 50g of standard lab food that was melted using a microwave. The sucrose was thoroughly mixed and dissolved completely before pouring 3 ml of food in standard vials. For all experiments fly crosses were maintained at 18°C on SF. Once adults eclosed, they were aged 3-5 days at 18°C and then shifted to 29°C for respective experimental regimes. For knockdown experiments flies were maintained at 29°C for 7 days on SF and then shifted to HSD (29°C) for another 7 days before collection. For over-expression experiments, flies were maintained at 29°C for 3-4 days and then transferred to HSD (29°C) for 4-5 days before collection. For all radioactivity feeding experiments the radioactive compounds were added in the HSD and feeding was started from day one of transfer to HSD (29°C). For lipid mobilization assays the flies were kept on [1-^14^C]-oleate containing HSD for 3 days and then transferred to cold SF for radioactive chase. For lipid incorporation assays the flies were transferred to [U-^14^C]-Sucrose containing HSD and samples were collected every 24 hours.

*Preparation of radioactive food:* 4 μCi of radioactive material was prepared in 15 μl Ethanol ([1-^14^C]-oleate) or water ([U-^14^C]-Sucrose) along with 5 μl of FIDC blue food dye (to visually confirm that the radioactive material is spread evenly on the food surface). HSD food was poured into empty vials without creating any bubbles to make sure to have an even surface when the food solidifies. The radioactive mixture was added on the surface of solidified HSD food dropwise using a pipette and making sure that it is evenly distributed. Importantly, the vials were then appended to a rotor that allowed slow rotation of the vials along the longitudinal axis to assist even spreading of the liquid as it dries overnight.

### BODIPY staining and imaging

The abdominal dorsal cuticle was dissected in relaxing buffer (1X PBS, 5 mM MgCl2 and 5 mM EGTA) using micro-scissors with the adipose tissue attached to the cuticle as described before (Rajan et al., 2017). The samples were fixed with 4% paraformaldehyde in relaxing buffer for 20 min. Subsequently the samples were washed with PBS, permeabilized with 0.1% PBT (PBS + 0.1 % TritonX100) in PBS for 30 min. PBT was removed by washing with PBS, 3 times, 5 min each, before adding BODIPY. For BODIPY staining, 500 μl of BODIPY in PBS (1/500 dilution of a 1 mg/ml stock in DMSO) was added to the samples and the samples were placed on a rotator for 30 min at room temperature. The samples were then washed in PBS, 2 times, 10 min each, incubated in PBS with DAPI for 10 min, washed 2 more times in PBS, 10 min each, and then mounted with Vectashield mounting media (VectaShield 1000). Samples were mounted with a bridge using two pieces of scotch tapes (3M) with the adipose tissue facing the cover slip. Samples were imaged using a Zeiss LSM780 confocal microscope. Images were acquired at room temperature.

### Triacylglycerol (TAG) assays

Whole animal TAG was measured as described previously (Tennessen et al., 2014). Briefly, 8 males were homogenized in 96 well deep well plates with 250 μl of ice cold PBT using a TissueLyser II (1 mm Zirconium beads, frequency: 30/sec, time: 30 sec). The plates were centrifuged at 1500 rpm for 3 min to remove any debris and 10 μl of the supernatant and triglyceride standards (Sigma: G7793-5ML) was added to 20 μl of triglyceride reagent (Sigma: T2449-10ML) in 96 well black clear bottom plates (Greiner bio-one; non-binding, black plates 655906) and the mixture was incubated at 37°C for 40-45 min. 100 μl of free-glycerol reagent (Sigma: F6428-40ML) was subsequently added to each well and the samples were incubated at 37°C for 5-10 min before reading absorbance at 540 nm using a 96 well plate reader.

### Immunohistochemistry to detect Pvf1 and PvR proteins

Rat anti-Pvf1 and Rat anti-PvR sera were used at a dilution of 1/200 and 1/500 respectively in blocking solution (BS: 1X PBS+0.1% TritonX100+5% BSA). Anti-Rat secondary antibody was used at a dilution of 1/500. Rabbit monoclonal antibody to p4EBP and secondary antibody (Donkey anti-Rabbit-488) for p4EBP detection were pre-cleared with fixed embryos (*w^1118^*) at a dilution of 1/50 in BS and then used at a dilution of 1/200 and 1/500 respectively in BS.

### Oenocytes/Adipose Tissue

The abdominal dorsal cuticle was dissected in relaxing buffer (1X PBS, 5 mM MgCl2 and 5 mM EGTA) using micro-scissors with the adipose tissue attached to the cuticle. The samples were fixed with 4% paraformaldehyde in relaxing buffer for 20 min. The samples were then washed in 1X PBS 3 times, 5 min each. The samples were permeabilized with 1X PBS+0.1% TritonX100 (PBT) for 30 min and then incubated with BS for 2 hours at room temperature. Primary antibody staining was performed in BS at 4°C overnight or 48 hours for staining the oenocytes with constant rotation (a long incubation is necessary for the antibody to percolate into the oenocytes) with constant rotation. Post primary incubation, samples were washed generously (5 times, 15-minute washes with PBT) at room temperature. Secondary, antibody incubation was done at room temperature for 2 hours in BS diluted 1/5 in PBT. Post secondary-incubation samples were washed in PBT and mounted with Vectashield mounting media. For phalloidin staining, PBT was exchanged with PBS with three washes and samples were incubated with phalloidin in PBS for 30 min at RT. Samples were then washed in PBS and mounted with vecta-shield mounting media.

### Muscle

Adult male thoraxes were prepared for fixation by removing the head and abdomen. Additionally, the tips of the legs were cut using micro-scissors to allow easy access to the leg muscles for the fixative. The dissected samples were fixed in 4% paraformaldehyde for 30 min. The samples were subsequently embedded in 4% low-melt agarose and left at 4 °C over night for the agarose to solidify. Samples in agarose blocks were mounted on to the stage of a vibratome (Leica VT1000M) in ice-cold PBS and 100 μm sections were cut. The sections were further trimmed under the microscope tissue sections with some surrounding agarose were transferred to 2 ml tubes. Subsequent immuno-staining steps were similar to as discussed in the previous section. Stained sections were mounted in Vectashield mounting media and imaged using a Zeiss LSM 780 confocal microscope.

### RT-qPCR

Real-time quantitative PCR (RT-qPCR) experiments were conducted using a Biorad CFX 96/384 device. The iQ SYBR green super mix and i-Script RT-reaction mix was used as per the manufacturer’s instructions for the qPCR reactions. 7.5 μl of the 2x reaction mix, primer-mix final concentration of 133 nM and 20 ng of cDNA (assuming 1:1 conversion of RNA to cDNA) were regularly used per reaction. Reagents and samples were dispensed using a Formulatrix Mantis small volume liquid handler. The ΔΔCt method was used to calculate fold change in experimental conditions. 4-5 biological replicates were used in all experiments. Transcript levels were normalized to *Drosophila Rp49*, *tub* and *gapdh*. Standard curves were run for each primer before use and we only used primers that showed an efficiency above 95%.

### Folch extraction and Thin-layer Chromatography (TLC)

#### Folch Extraction

Animals were collected in 1.5 ml screw cap tubes along with 1 mm zirconium beads and were frozen on dry ice. Samples were stored at −80°C if necessary, until all samples were ready for processing. For processing of the samples, 600 μl of Methanol:ChCl3:H2O (10:5:4) was added to the tubes using a graduated glass Hamilton syringe and the animals were homogenized using the Qiagen TissueLyser II instrument. Two rounds of homogenization at a frequency of 30/sec and total duration of 30 sec per round were performed to ensure complete homogenization of the tissues. The samples were placed on a rotor at 37°C for 1 hour. Subsequently, 160 μl of ChCl3 and 160 μl of 1 M KCl were added to each sample. The samples were shaken vigorously for 5 sec and briefly vortexed before centrifuging at 3000 rpm for 2 min on a standard Eppendorf bench top centrifuge for phase separation. 220 μl of the organic phase (bottom layer) was pipetted out into new clean 1.5 ml centrifuge tubes using a graduated glass Hamilton syringe. The samples were then placed in a vacuum concentrator (Labcono centrivap console) and the organic solvent was completely removed. The dried samples were stored at −80 if needed or run directly on a TLC plate.

#### Thin-Layer Chromatography

For TLC, the dried samples were resuspended in 40 μl CHCl3 and the entire volume was loaded on to Analtech channeled TLC plates with preadsorbent zones that allow loading of large volumes of samples (Analtech P43911). The lipids were then separated using hexane:diethyl ether:acetic acid (80:20:1) solvent system. The plates were exposed to phosphor imager screens overnight and revealed by using a Typhoon FLA 7000 phosphor imager. The density of the TAG bands was calculated using imageJ/Fiji.

### Preparation of single nuclei suspension from adult *Drosophila* abdominal cuticle

40 adult male abdomens were quickly dissected in ice cold relaxing buffer (1xPBS, 10 mM EGTA and 10 mM MgCl2) by cutting off the last 2^nd^ and abdominal segment with micro-scissors and hollowing out the abdomen by removing the intestines and male reproductive organs. The dissected tissue was then roughly chopped and transferred to a Dounce homogenizer. The relaxing buffer was replaced with 1.3 ml of nuclei lysis buffer from the Sigma NUC-201 Nuclei isolation kit. Single nuclei suspension from the homogenate was prepared according to the manufacturer’s instruction the in the NUC-201 kit.

Briefly, a 1.5 M sucrose cushion was used for generating the gradient for density gradient centrifugation. Density gradient centrifugation was performed in 2 ml tubes using a SW55Ti rotor on a Beckman ultracentrifuge.

### Analysis of snRNA-Seq data

We used the 10X Genomics cellranger count pipeline (version 3.1.0) to analyze the demultiplexed FASTQ data and generated the single cell count matrix, once for each sample. We aligned the reads to a custom “pre-mRNA” reference which was generated as described by 10X Genomics based on FlyBase R6.29. We applied SoupX (version 0.3.1) (Young and Behjati, 2018) to directly correct the count matrix from cellranger with fixed contamination value equals 0.45 for each sample. We filtered the cells beyond UMI counts ± 2-fold Standard Deviation of the average total sample counts (log10) after SoupX, which were regarded as doublets or dead cells in droplet. The quality filtered datasets were combined into a single Seurat (version 3.1.2) object (Stuart et al., 2019) and integrated using Harmony (version 1.0) (Korsunsky et al., 2019) with default analysis workflow and parameters. A resolution of 0.1 was chosen as clustering parameter. The code for the snRNA-Seq analysis can be found at https://github.com/liuyifang/Drosophila-PDGF-VEGF-signaling-from-muscles-to-hepatocyte-like-cells-protects-against-obesity. Dot and Violin plots were generated using the Seurat DotPlot and VlnPlot functions. We performed pathway enrichment analysis on marker genes with positive fold change for each cluster using a program written in-house. Gene sets of Transcription Factor (TF) target genes of major signaling pathways were assembled manually (unpublished data). Enrichment P-value was calculated based on the hypergeometric distribution using the background of 11863 genes identified as expressed in this dataset. The strength of enrichment was calculated as negative of log10(p-value), which is used to plot the heatmap.

### Quantification and Statistical analysis

Graphical representation and statistical analysis of all quantitative data was performed using GraphPad Prism 8 software (www.graphpad.com). Quantification of lipid droplet size was performed using CellProfiler and the pipeline used will be made available upon request to the corresponding author (Lamprecht et al., 2007). Quantification of fluorescent intensities of immunostained samples was performed using a custom-made ImageJ (Fiji) macro (also available upon request).

For starvation resistance studies, the data is presented as Kaplan-Meier survival plots. Statistical significance for the survival plots was determined using the Log-Rank (Mantel-Cox) test. For all other quantifications, including the data for TAG quantification, the data is presented as either bar plots or dot plots with the error bars showing standard error of mean. Methodologies used for determining statistical significance are mentioned in the figure legends. Asterisks illustrate statistically significant differences between conditions. ****p<0.0001, 0.0001<***p<0.001, 0.001<**p<0.01 and 0.01<*p<0.05

---

**Figure S1.**
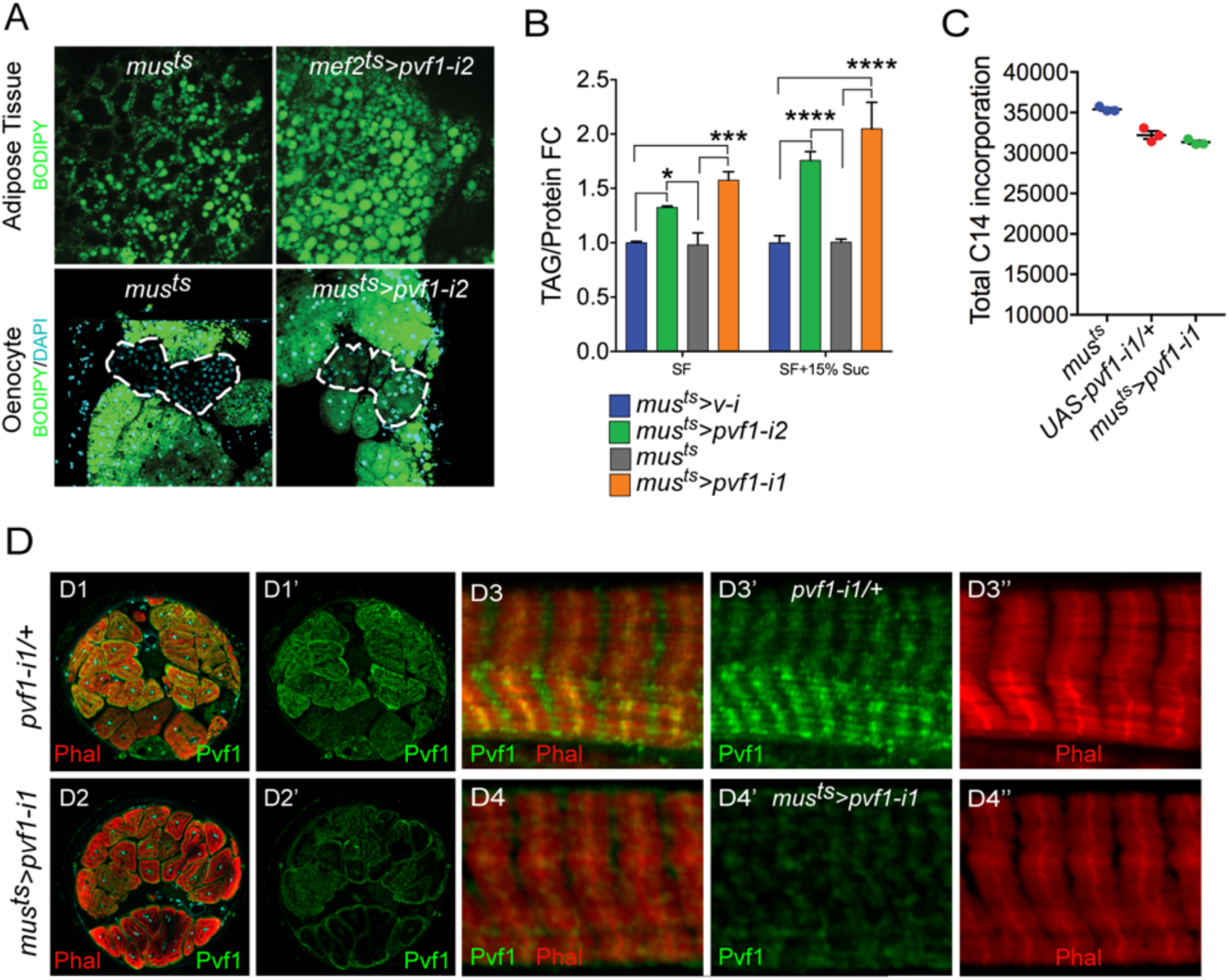
(Related to Figure 1): A muscle to oenocyte Pvf1 signaling axis protects against obesity (A) BODIPY staining showing neutral lipid accumulation in the adult male adipose tissue (AT) and oenocytes (dorsal abdominal cuticle) of control (*mus^ts^= muscle-Gal4^Gal80ts^*) flies and flies with muscle-specific knockdown of *pvf1* (*mus^ts^>pvf1-i2*) (NIG: 7103R-2) (B) Total TAG content of adult males with adult muscle-specific knockdown of *pvf1* (*mus^ts^>pvf1-i2*) (NIG: 7103R-2) reared on standard lab food (SF) and 15% w/v High-sugar diet (HSD). (SF = standard lab food). (SF+15% Suc = standard lab food supplemented with 15% sucrose w/w). n=6, Two-Way ANOVA followed by Tukey’s HSD test. (C) ^14^C counts per minute of whole fly homogenates of control (*mus^ts^*) and *UAS-pvf1-i1/+* flies and flies with muscle-specific knockdown of *pvf1* (*mus^ts^>pvf1-i1*) (VDRC: kk102699) that were fed with [U-^14^C]-Glucose for 48 hours. n=3, One-Way ANOVA followed by Tukey’s HSD test. (D) Distribution of Pvf1 protein (anti-Pvf1 antibody:: Green) in the leg muscle of control (*pvf1-i1/+*) (D1, D1’, D3, D3’) flies and flies with muscle-specific knockdown of *pvf1* (*mus^ts^>pvf1-i1*) (VDRC: kk102699) (D2, D2’, D4, D4’). Figures D1, D1’ and D2, D2’ show cross-section of the leg muscle. Figures D3-D3’ and D4-D4’ show transverse section of the leg muscle.

**Figure S2.**
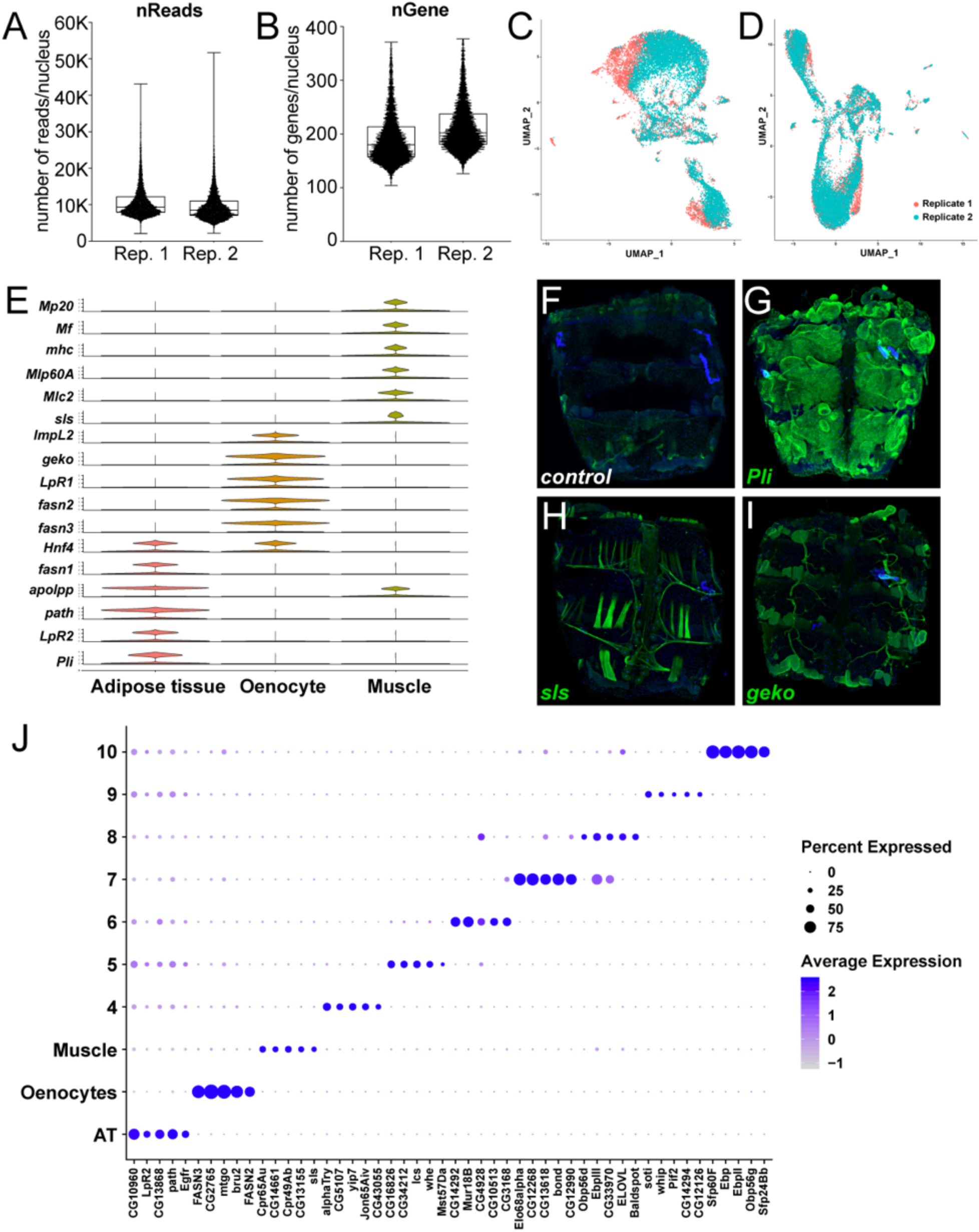
(Related to Figure 2): snRNA-seq of adult fly abdomens: Validation of marker genes and top marker genes per cluster (A) Distribution of number of reads per nucleus in each replicate (Rep. 1 and 2). (B) Distribution of number of genes per nucleus in each replicate (Rep. 1 and 2). (C) UMAP plots representing batching due to replicates prior to Harmony (D) UMAP plots representing correction of the batch effects after Harmony. (E) Violin plots representing the normalized expression of marker genes specific for the metabolically active tissues: adipose tissue, oenocytes, and muscle. (F-I) Validation of novel marker genes for the three major clusters Adipose tissue, muscle and oenocyte. Trojan-Gal4 lines for each of the candidate genes (*pli^MI00302-TG4.2^*, *sls^MI10783-TG4.1^* and *geko^MI02663-TG4.1^*) were used to drive expression of EGFP to mimic the pattern of expression of the genes themselves. One week old adult male abdomens were dissected and stained with anti-EGFP to revel the expression pattern of each of the three genes. (F) *w^1118^* adult male abdomen stained with anti-EGFP shows background staining pattern. Muscle cells that were damaged during dissection tend to uptake the antibody and show some staining on the edges of the sample. (G) *pli^MI00302-TG4.2^>UAS-2xEGFP* adult male abdomens show EGFP staining primarily in the adipose tissue. Staining was not seen in the abdominal muscles or in the oenocytes. (H) *sls^MI10783-TG4.1^>UAS-2xEGFP* adult male abdomens show EGFP staining in the abdominal tubular muscles and in the alary muscles. (I) *geko^MI02663-TG4.1^>2xEGFP* adult males abdomens show EGFP staining in the oenocytes and in the tracheal tubes. Staining above the background sample was not seen in the adipose tissue or the abdominal muscles. (J) Dot plot representation of top 5 genes enriched per cluster based on average logFC. The size of the dot represents the percentage of cells expressing a gene while the color gradient represents the level of gene expression.

**Figure S3.**
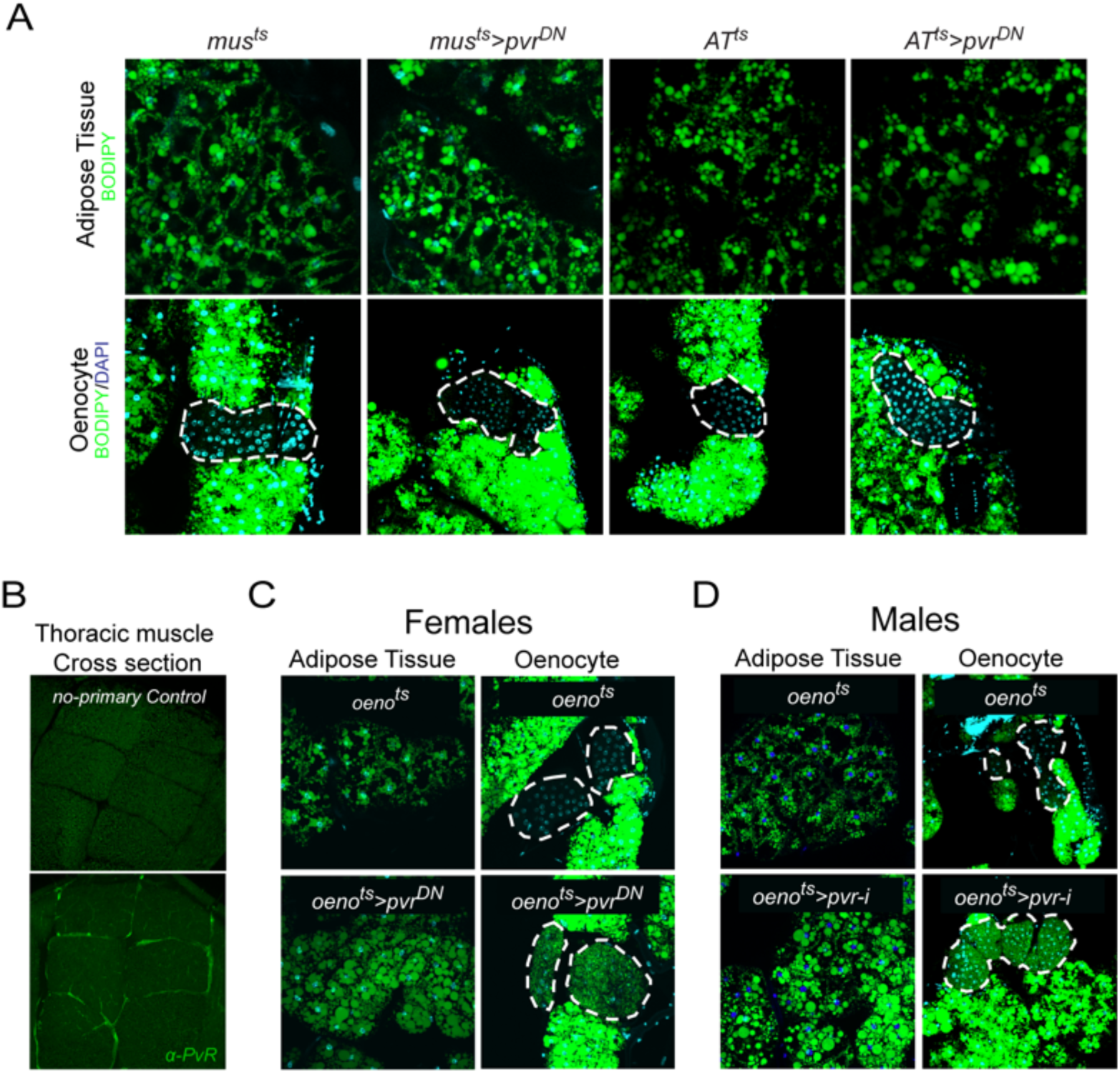
(Related to Figure 2): PvR signaling works specifically in the oenocytes to protect against obesity (A) BODIPY staining showing neutral lipid accumulation in adult male adipose tissue (AT) and oenocytes (dorsal abdominal cuticle) of control males and males over-expressing *pvr^DN^* in the adult muscle (*mus^ts^>pvr^DN^*) and adipose tissue (*AT^ts^>pvr^DN^*). Loss of PvR signaling in the muscle or adipose tissue does not have any effect on the neutral lipid content of either the adipose tissue or the oenocytes. (B) Distribution of PvR protein (anti-PvR antibody::Green) in the thoracic muscle of control (*mus^ts^*) males. A cross-section of the indirect flight muscles is shown. (C) BODIPY staining showing neutral lipid accumulation in the adult female adipose tissue and oenocytes (dorsal abdominal cuticle) of control (*oeno^ts^=oenocyte-Gal4^Gal80ts^*) flies and flies with oenocyte-specific over-expression of (*oeno^ts^*>*pvr^DN^*). (D) BODIPY staining showing neutral lipid accumulation in the adult male adipose tissue and oenocytes (dorsal abdominal cuticle) of control (*oeno^ts^*) flies and flies with oenocyte-specific knockdown of *pvr* (*oeno^ts^*>*pvr-i*) (NIG: 8222R-2).

**Figure S4.**
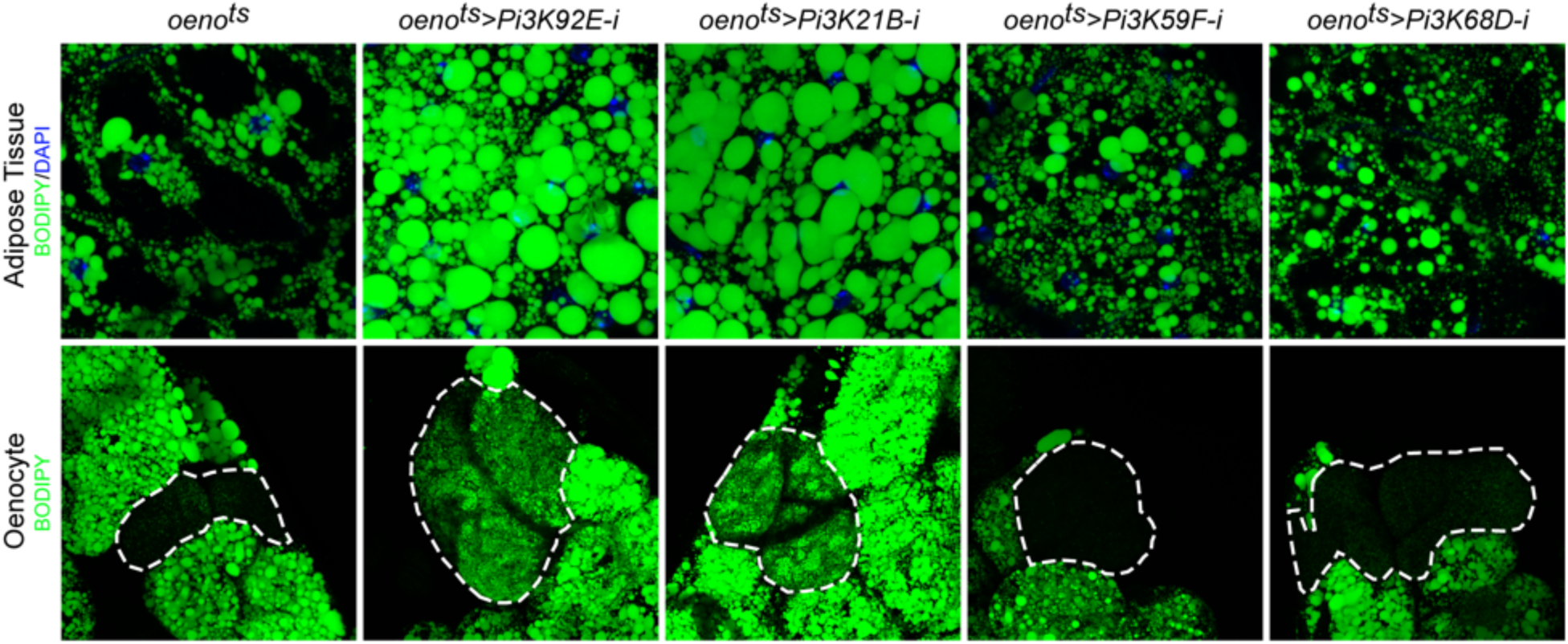
(Related to Figure 3): Oenocyte-specific loss of only Pi3K92E and Pi3K21B leads to obesity BODIPY staining showing neutral lipid accumulation in the adult male adipose tissue and oenocytes (dorsal abdominal cuticle) of control (*oeno^ts^=oenocyte-Gal4^Gal80ts^*) flies and flies with oenocyte-specific (*oeno^ts^>*) knockdown of: *Pi3K92E; Pi3K21B, Pi3K59F* and *Pi3K68D*.

**Figure S5.**
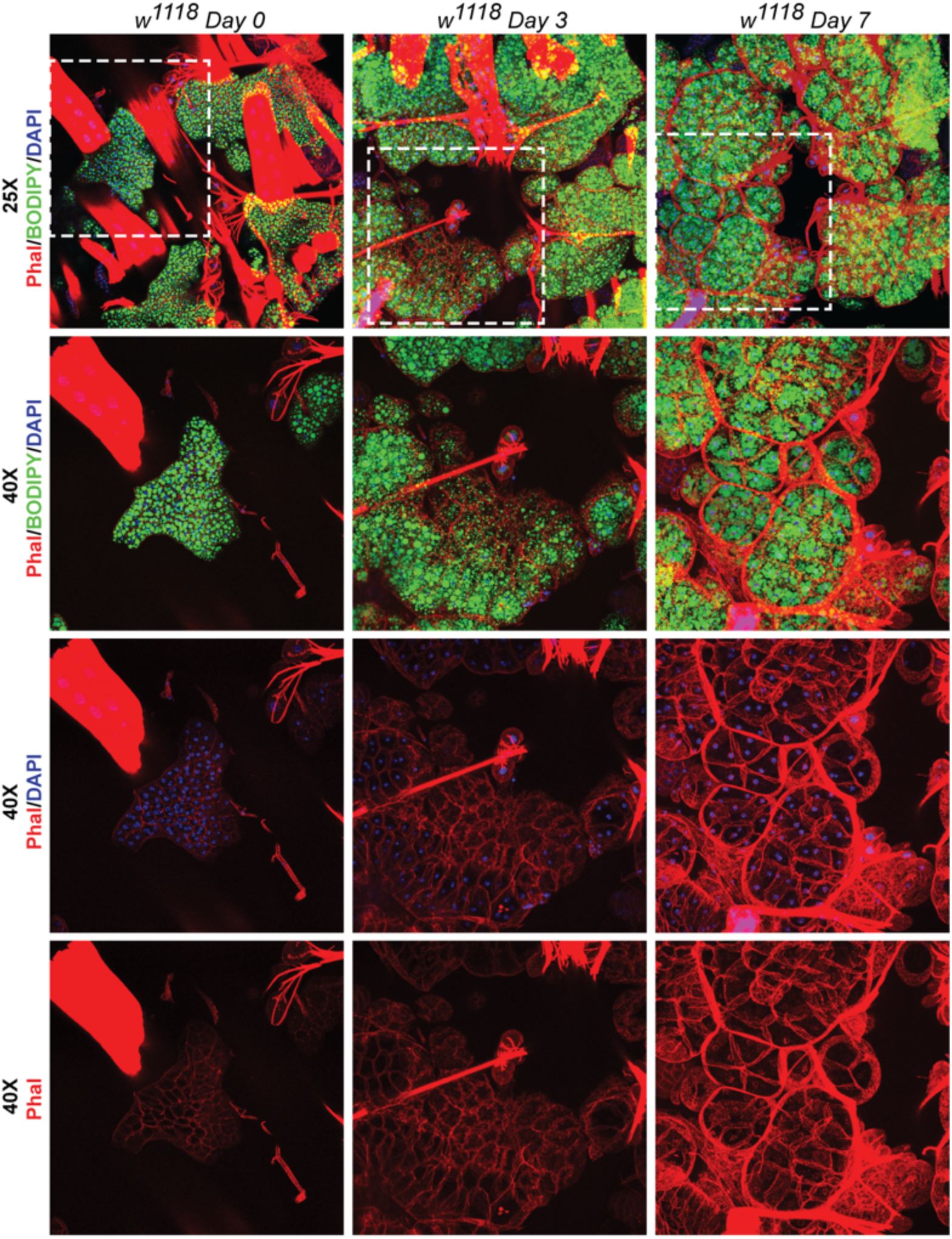
(Related to Figure 6): Recovery of adipose tissue lipid stores involve formation and expansion of new lipid droplets, and, expansion of the adipose tissue cells BODIPY, DAPI and Phalloidin staining of adipose tissue preparations from *w^1118^* flies on day0, day3 and day7 after eclosion. Topmost row shows images acquired with a 25x eyepiece. Bottom three rows show images acquired with a 40x eyepiece. Dotted white boxes show the regions that were re-imaged with the 40x lens. On day 0 post-eclosion the adipose tissue is small with each cell having two nuclei that are positioned very close to each other. By day 3 and day 7 the adipose tissue cells become progressively larger and fill up with more lipid droplets per cell (both large and small).

**Figure S6:**
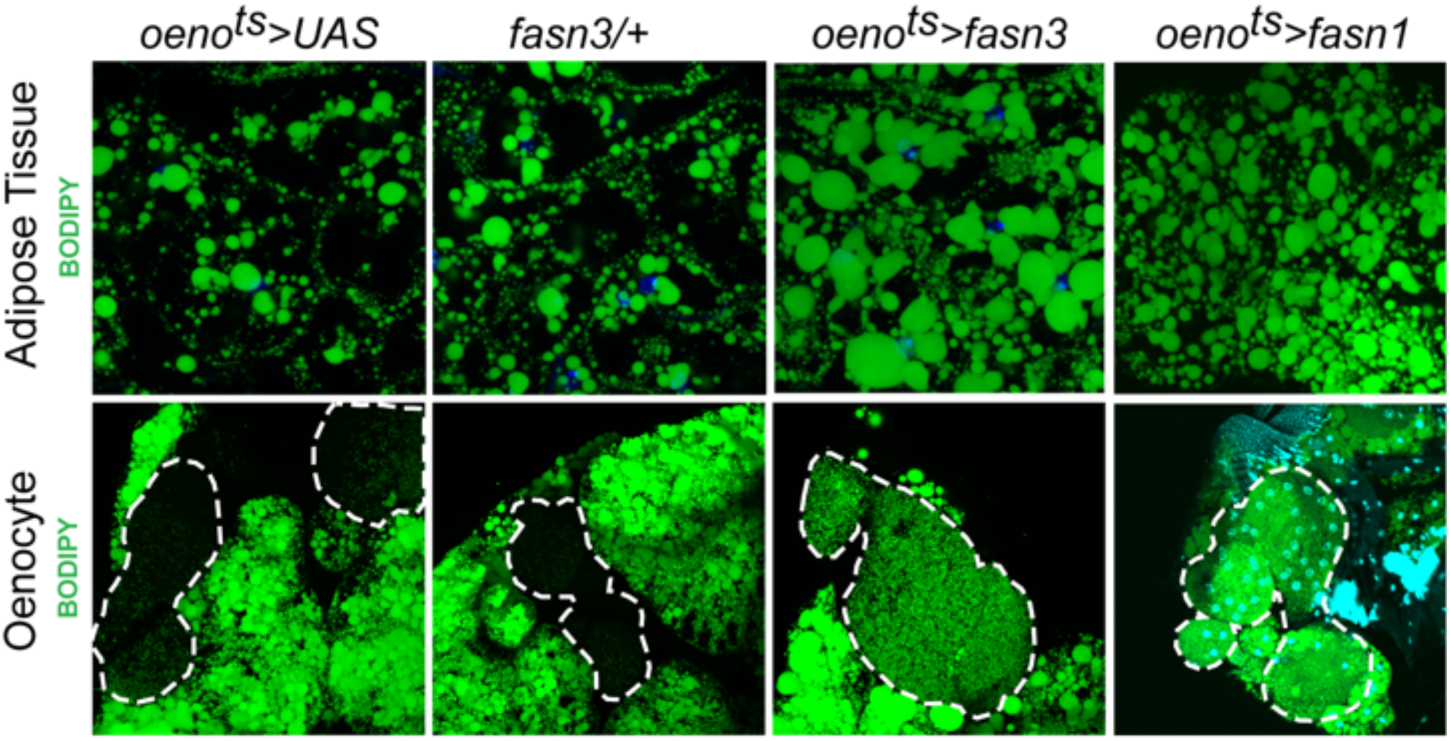
Oenocyte-specific activation of lipid synthesis leads to increased lipid stores in the adipose tissue BODIPY staining showing neutral lipid accumulation in the adult male adipose tissue and oenocytes (dorsal abdominal cuticle) of control (*oeno^ts^>UAS*) flies and flies with oenocyte-specific over-expression of *fasn3* and *fasn1* (*oeno^ts^>fasn3* and *oeno^ts^>fasn1*). *oeno^ts^=oenocyte-Gal4^Gal80ts^*

